# Systematic Evaluation Defines the Limits of Ferroptosis in Cancer Therapy

**DOI:** 10.64898/2026.03.11.711115

**Authors:** Kenji M. Fujihara, Azmi Aziz, Behnia Akbari, Maria Gutierrez-Perez, Gregory Francis, Siham Zentout, Kameng Wu, Nicholas J. Clemons, Erdem M. Terzi, Michael E. Pacold, Richard Possemato

**Affiliations:** Department of Pathology, New York University Grossman School of Medicine, New York, NY 10016, USA; Laura & Isaac Perlmutter Cancer Center, New York, NY 10016, USA; Department of Radiation Oncology, New York University Grossman School of Medicine, New York, NY 10016, USA; H12O-CNIO Lung Cancer Clinical Research Unit, Instituto de Investigación Hospital 12 de Octubre (i+12) & Centro Nacional de Investigaciones Oncológicas (CNIO), Madrid, Spain; Department of Biochemistry and Molecular Pharmacology, New York University Grossman School of Medicine, New York, NY, USA; Division of Cancer Research, Peter MacCallum Cancer Centre, Parkville, Victoria, Australia. Peter MacCallum Department of Oncology, University of Melbourne, Parkville, Victoria, Australia

## Abstract

Ferroptosis is a cell death mechanism characterized by the accumulation of iron-catalyzed lipid peroxides in membrane lipid acyl chains and subsequent loss of membrane integrity.^1^ Despite thorough investigation of its mechanisms in cultured cells, induction of ferroptosis has unresolved clinical utility in cancer therapy. Here, we systematically evaluate ferroptosis induction via multiple mechanisms, in both cell and tumor models, using focused genetic screens, genetic loss-of-function systems, and pharmacological perturbations. Through this analysis we identify cancer cell line subsets with distinct responses to canonical ferroptosis inducers and suppressors and define the underpinnings of each. Inhibition of central *in vitro* ferroptosis suppressors GPX4, GCLC, or SLC7A11 across these multiple models fails to impact established tumor growth. In contrast, deficiency in the cytosolic thioredoxin reductase and pharmacologic GCLC inhibition potently induces tumor regression and triggers a form of non-ferroptotic cell death regulated by cystine availability and translation. These analyses further reveal that the principal essential function of environmental cystine in cultured cells is to support selenoprotein function, identified through investigating our finding that β-mercaptoethanol supports exponential growth in cystine-free conditions. Thus, while ferroptosis activation may be efficacious alone or in combination with other therapies in specific tumor contexts, cell culture systems greatly overestimate the potential anti-cancer effects of ferroptosis induction via the GPX4 axis.

## Main

The study of ferroptosis can be traced back to the first investigations of amino acid requirements in mammalian cell culture systems, as cystine-deprivation potently induced this cell death mechanism.^2^ The first pharmacological ferroptosis inducers (erastin and RSL-3) originated from high-throughput phenotypic drug screens aimed at identifying genotype-selective anti-cancer agents in oncogenically transformed HRAS mutant fibroblasts.^3,4^ Erastin blocks cystine uptake by inhibiting the cystine-glutamate antiporter (system xCT) component SLC7A11.^5^ RSL-3 covalently modifies and inhibits the glutathione (GSH)-dependent peroxidase and selenoprotein GPX4,^4,6^ which repairs lipid peroxides generated by oxidative damage to membrane lipid polyunsaturated fatty acid (PUFA) acyl chains.^7,8^ Depleting GSH through cystine deprivation or inhibition of GSH synthesis also limits GPX4 function, facilitating unchecked PUFA oxidation.^3,6^ Alteration of intracellular free iron levels or membrane lipid content potently modulates ferroptosis activation: iron availability toggles the degree of ongoing oxidative damage, while high PUFA content provides extra substrate for lipid peroxidation.^3,6,9^ In cultured cells ferroptosis can be suppressed by lipid radical trapping antioxidants (RTA).^3^ In particular, CoQ_10_ is a key cellular RTA limiting ferroptosis and is recycled in lipid membranes by the oxidoreductase FSP1,^10,11^ while tetrahydrobiopterin,^12^ squalene,^13^ and cholesterol or its derivatives^14–16^ limit ferroptosis in specific contexts. Exogenously supplied synthetic or natural lipophilic RTAs, such as Ferrostatin-1 (Fer-1), Liproxstatin-1 (Lip-1) and vitamin E (VitE), block both cystine-deprivation induced or GPX4-inhibition induced ferroptosis.^3,6^

While studies have identified cell-state-dependent ferroptosis modifiers,^17^ much work in the field has focused on a small number of cell lines, including those that are highly susceptible to ferroptosis. As a result, seemingly contradictory results have arisen, most notably the relative inability of GSH synthesis inhibition to potently induce ferroptosis compared to GPX4 inhibition. Moreover, genetic screens to identify novel ferroptosis regulators frequently arrive at highly context-specific results.^18^ Thus, leveraging ferroptosis activation as a promising avenue for anti-cancer therapy has remained enigmatic.

## CRISPR tiling mutagenesis provides target validation of ferroptosis inducers

To begin to address these questions, we perform CRISPR-based mutagenesis of key genes (including *GPX4*, *GCLC*, and *SLC7A11*) to verify the specificity and targets of compounds commonly utilized to study ferroptosis: GPX4 inhibitor RSL-3, SLC7A11 inhibitor erastin, and GCLC inhibitor BSO (**Figure 1A**). To maximize gene coverage we utilize the enhanced AsCas12a variant (enAsCas12a), which harbors a flexible PAM, in addition to SpCas9.^19^ HT-1080 fibrosarcoma cells transduced with CRISPR tiling libraries were exposed continuously to lethal doses of these compounds (**Extended Data Figure 1A**), and CRISPR RNAs (crRNAs) or single guide RNAs (sgRNAs) driving resistance identified (**Figure 1B-C, Extended Data Figure 1B, Supplementary Table 1**). Screens utilizing enAsCas12a better enriched for crRNAs targeting specific gene regions, while those utilizing SpCas9 exhibited superior negative selection for sgRNAs targeting essential genes. Therefore, we focus on crRNAs enriched in the enAsCas12a screen hereafter.

**Figure 1.**
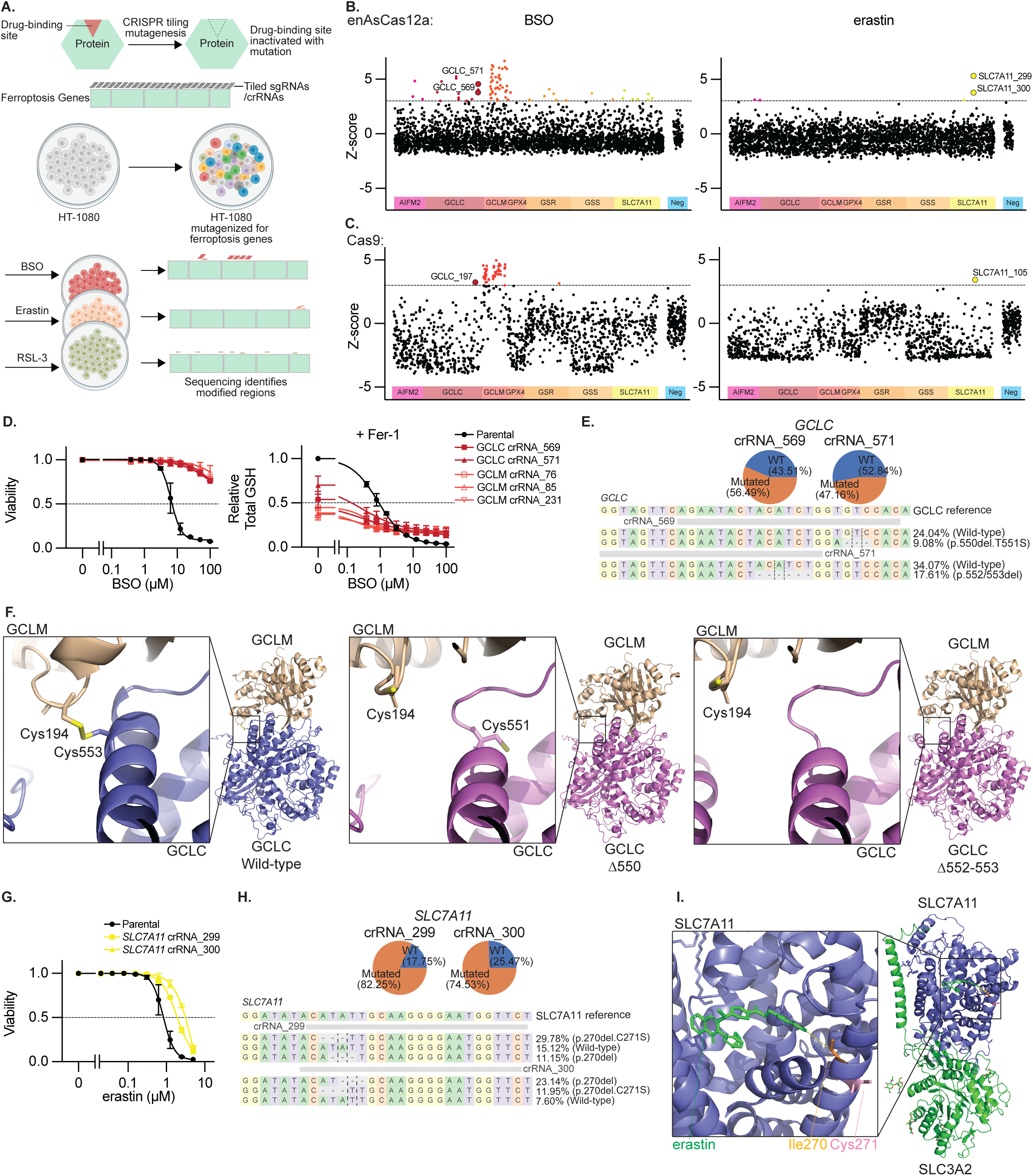
CRISPR tiling mutagenesis provides target validation of ferroptosis inducers. **A**, Schematic outlining tiling mutagenesis approach. **B-C**, Z-score of guide RNA enrichment using enAsCas12a (B) or Cas9 (C), tiled across genes indicated upon BSO (100μM) or erastin (5μM) selection, dotted line at Z=3, n=2 biological replicates. **D**, Viability (ATP, left) and GSH content (right) of HT-1080 cells harboring enAsCas12a and indicated guides upon BSO or Fer-1 (10 μM) treatment, 3 days. Data are mean ± SEM, n=3 biological replicates. **E**, Proportion of mutant or WT *GCLC* alleles upon enAsCas12a and indicated guide RNA transduction (top). Guide RNA targeting region and major mutant allele for each shown (bottom). **F**, AlphaFold 3.0 Model of GCLM:GCLC binding for WT GCLC (left) and mutants identified (center and right). Insets show targeted region. **G**, Viability (ATP) of HT-1080 cells harboring enAsCas12a and indicated guides upon erastin treatment, 3 days. Data are mean ± SEM, n=3 biological replicates. **H**, Proportion of mutant or WT *SLC7A11* alleles upon enAsCas12a and indicated guide RNA transduction (top). Guide RNA targeting region and major mutant alleles for each shown (bottom). **I**, SLC7A11:SLC3A2 binding to erastin (green) from PDB (7EPZ)^23^. Inset shows targeted region and residues modified (I270-yellow, C271-pink).

Those crRNAs enriched upon BSO treatment target distinct regions in *GCLC* or broadly target *GCLM,* whose loss is known to cause BSO resistance,^20,21^ and we validated several of these crRNAs (**Figure 1B, Extended Data Figure 1C-D**). Mutagenized and selected populations were completely resistant to the toxic effects of BSO up to 100 μM (**Figure 1D**), and upon BSO treatment retain GSH at 10% of wildtype levels (**Figure 1D**), similar to what others have observed upon *GCLM* disruption.^21^ Sequencing of the *GCLC* crRNA targeting sites revealed that approximately half of the reads are mutated, implying that a single mutant allele is sufficient to drive resistance (**Figure 1E**). The most common mutations map to a region essential for GCLC: GCLM heterodimer formation, GCLM residue C194 and GCLC residue C553, disrupting a disulfide bond (**Figure 1F**)^22^. Overexpression of a GCLC cDNA or cDNAs harboring positively selected GCLC mutants induce resistance to BSO (**Extended Data Figure 1E**).

The crRNAs enriched upon erastin treatment (*SLC7A11* cr299, cr300), induce significant but not complete resistance to erastin-induced ferroptosis (**Figure 1G**, **Extended Data Figure 1D**). Sequencing of positively selected cells revealed that ∼75% of reads were mutated, producing the same two dominant mutant alleles (**Figure 1H**). These crRNAs target SLC7A11 residue I270, located in a transmembrane helix that forms part of a hydrophobic cluster bound by erastin (**Figure 1I**)^23^. Overexpression of a wild-type SLC7A11 cDNA or those harboring positively selected SLC7A11 mutants induced resistance to erastin (**Extended Data Figure 1F**).

No crRNAs or sgRNAs were significantly enriched in cells treated with RSL-3, as well as two additional GPX4 inhibitors (ML-162 and ML-210), although resistance to ML-210 was observed in the presence of the mutagenesis library (**Extended Data Figure 1G-I**). Overexpressing GPX4 induced RSL-3 partial resistance, and a selenocysteine to cysteine mutant (GPX4^U46C^)^24,25^, which retains partial GPX4 catalytic activity but cannot interact with RSL-3, induced complete resistance to RSL-3, confirming RSL-3 selectivity (**Extended Data Figure 1J**). Meanwhile GPX4 cDNA without the selenocysteine insertion sequence (SECIS), required for synthesis of the catalytic site selenocysteine, failed to rescue.

These results support prior results that BSO depletes cellular GSH and triggers ferroptosis through targeting GCLC,^26^ erastin triggers ferroptosis through targeting SLC7A11,^3^ and ferroptosis induction by RSL-3 requires GPX4 targeting,^4^ and provide additional tools for investigating induction of ferroptosis, as utilized below.

## Perturbations Inducing Ferroptosis *In Vitro* Fail to Trigger Tumor Regression

Having confirmed the relative selectivity of commonly used ferroptosis inducers for their targets, we evaluated the efficiency with which ferroptosis could be induced in cell culture and trigger tumor regression by performing targeted CRISPR/Cas9 drop-out screens and generating genetic models in which key ferroptosis targets could be inducibly suppressed. We generated a pool of sgRNAs comprised of ferroptosis-related genes (*AIFM2*, *GCLC*, *GCLM*, *GPX4*, *GSR*, *GSS*, *SLC7A11*) and known essential genes and cancer targets (*DBR1*, *EEF2*, *GAPDH*, *KIF11*, *PCNA*, *PLK1*, *POLR2B*, *PSMB1*, *RPA3*, *RPL3*), and transduced two cancer cell lines (CCLs: A549 and HT-1080), which vary in their ferroptosis sensitivity and GSH dependence (**Figure 2A**). We used loss of sgRNA abundance to infer gene requirement, rescue with Fer-1 to infer cell death by ferroptosis, and β-mercaptoethanol (βMe) to enable SLC7A11-indenpdent cystine uptake (**Figure 2B, Supplementary Table 2**). Of the ferroptosis-related genes, only *GPX4* sgRNAs were strongly depleted under standard culture conditions, which was rescued by Fer-1 but not βMe (**Figure 2B**). Surprisingly, sgRNAs targeting GPX4 were not depleted in tumor xenografts, and no other sgRNAs targeting ferroptosis-related genes were significantly depleted under any condition (**Figure 2B**). In contrast, all essential gene sgRNAs were depleted under every tissue culture condition and in tumor xenografts (**Figure 2B**). To ask whether these findings were related to insufficient sgRNA reagent coverage, we repeated these experiments using our SpCas9 tiling library targeting these genes, which yielded concordant results (**Figure 2C**, **Extended Data Figure 2, Supplementary Table 3**). sgRNAs directed against the signal sequence of the mitochondrial GPX4 isoform were not depleted (**Figure 2C**), consistent with the canonical view^27^ that cytoplasmic GPX4 activity is key to cell viability *in vitro*.

**Figure 2.**
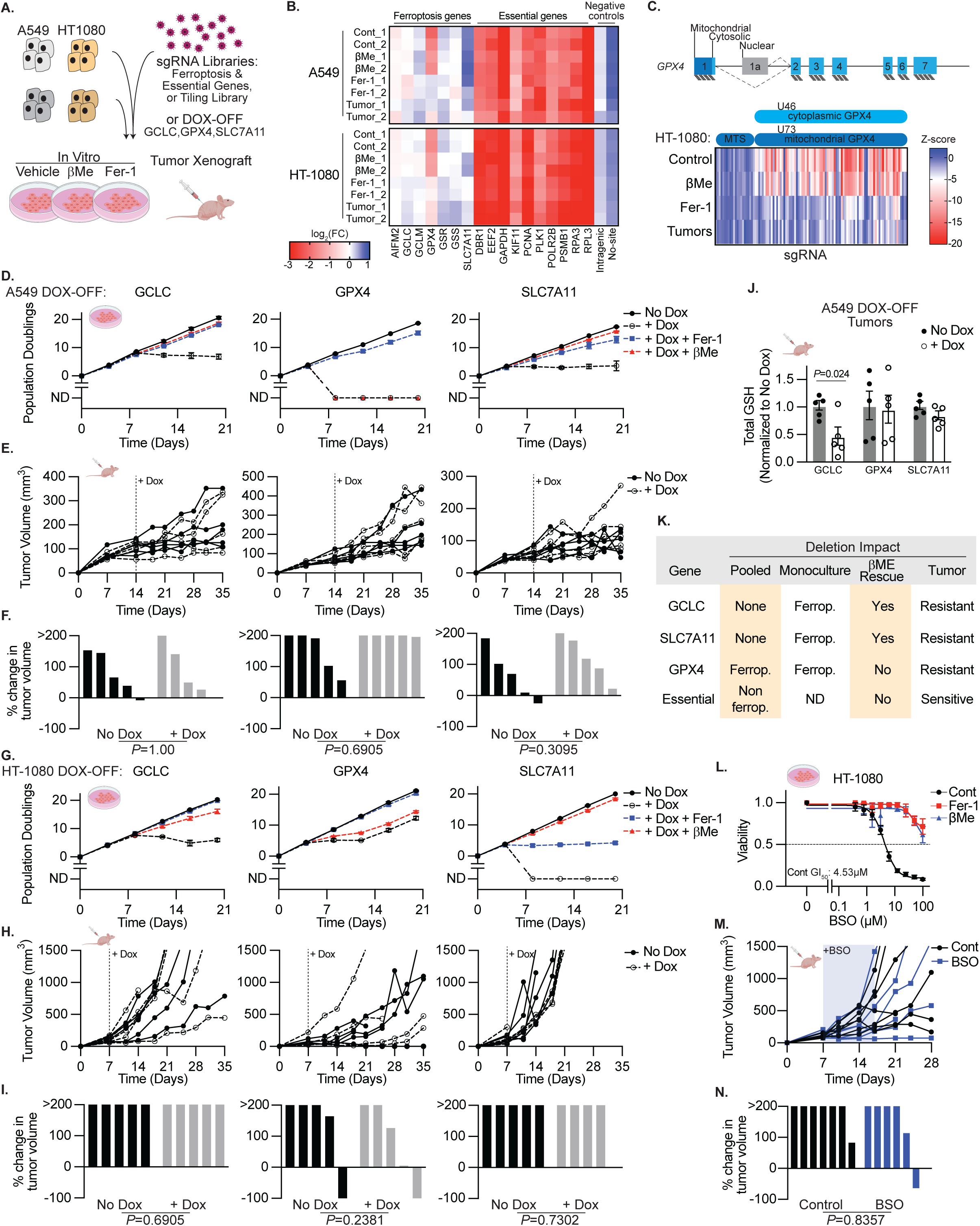
Perturbations Inducing Ferroptosis In Vitro Fail to Trigger Tumor Regression. **A**, Genetic screen design. **B**, Heatmap reporting log2 fold change in the abundance of sgRNAs targeting indicated genes or controls upon 15 days *in vitro* culture, including βMe (50 μM) or Fer-1 (10 μM) treatment, or upon growth as subcutaneous tumor xenografts, HT-1080 (14 days) or A549 (21 days) cells. Data are mean of n=10 guide RNAs, biological replicates reported separately. **C**, Schematic of *GPX4* genomic structure showing alternative splicing. Below, heatmap reporting Z-score of the abundance of sgRNAs targeting indicated regions within *GPX4* in HT-1080 cells grown in conditions from *A*. Data report mean of two biological replicates. **D,G**, Cumulative population doublings of A549 (*D*) or HT-1080 (*G*) cells harboring cDNAs encoding the indicated genes under doxycycline control with deletion of the corresponding endogenous gene. Cells cultured in the presence or absence of doxycycline (1 μg/mL), βMe (50 μM), or Fer-1 (10 μM). ND, no remaining detectable cells. Data report mean ± SD of n=3 technical replicates. **E,F,H,I**, Growth of individual subcutaneous tumors derived from cells in D-G with or without doxycycline supplemented food (1 g/kg, shaded region) reported as tumor volume (*E, H*) or change in tumor volume 21 days after doxycycline addition or ethical endpoint if reached earlier (*F, I*). Data are individual tumor measurements. P value reports result of Mann-Whitney U test. **J**, Total GSH levels in lysates of tumors from *G-H* at endpoint. Data report mean ± SEM of n=5 biological replicates. **K**, Summary of the impact of ferroptosis target gene deletion on cell viability and tumor growth. **L**, Viability (ATP) of HT-1080 cells treated with BSO, with or without βMe (50 μM), or Fer-1 (10 μM), 3 days. Data report mean ± SEM of n=3 biological replicates. GI50 indicated. **M-N**, Growth of individual subcutaneous tumors derived from cell lines in *A* with or without BSO in drinking water (shaded region) reported as tumor volume, (*M*), or change in tumor volume 21 days after start of BSO treatment or ethical endpoint if reached earlier (*N*). Data are individual tumor measurements. P value reports result of Mann-Whitney U test.

To use an orthogonal approach that would allow for inducible, near-complete suppression of each ferroptosis target, we generated single-cell CRISPR knockout (KO) clones of GPX4, GCLC and SLC7A11 with addition of doxycycline (DOX) suppressible cDNAs of each gene in HT-1080 and A549 cells (**Extended Data Figure 3 A-B**). Similar DOX-OFF models are highly responsive to low concentrations of DOX and can readily suppress the growth of cultured cells and tumors.^28,29^ We assessed the long term impact of gene suppression *in vitro*, with Fer-1 or βMe supplementation, and in tumor xenografts (**Figure 2D-I**). DOX suppressed GPX4, GCLC, and SLC7A11 protein levels (**Extended Data Figure 3 A-B**), leading to a complete suppression of in vitro cell proliferation (**Figure 2D,G**), with the caveat that GPX4 de-repression was observed in HT-1080 cells at extended timepoints. Loss of cell viability was blocked by Fer-1, confirming induction of ferroptosis (**Figure 2D,G**). The observed dependance on loss of SLC7A11 and GCLC differs from that observed in the pooled setting, as non-perturbed cells likely compensate for suppression of SLC7A11 or GCLC, but not GPX4 (**Figure 2B**). Consistent with the genetic screen results, suppression of GPX4, GCLC, or SLC7A11 in established tumors failed to impact tumor growth rates (**Figure 2E,F,H,I**), and tumors in which GCLC was suppressed had significantly lower levels of total GSH (**Figure 2J**). These data demonstrate that models in which ferroptosis can be readily and potently induced *in vitro* by deletion of SLC7A11, GCLC, and GPX4, fail to suppress the growth of established tumors (**Figure 2K**).

To further probe the discordance between these *in vitro* and tumor xenograft results and considering that those approaches do not systemically inhibit these targets, we utilized a pharmacological approach. BSO is orally available and has known pharmacokinetics appropriate for use *in vivo*.^30^ HT-1080 cells are sensitive to BSO *in vitro* (GI50 4.53 μM), which can be rescued by Fer-1 treatment, supporting ferroptosis induction (**Figure 2L**). However, established HT-1080 tumor xenografts failed to respond to BSO treatment, despite this BSO treatment regimen being the maximum tolerable (**Figure 2M-N**). Therefore, we conclude that *in vitro* models substantially overestimate the anti-cancer efficacy of ferroptosis induction.

## Phenotypic Modulatory Profiling Defines Therapeutic Index of *In Vitro* Ferroptosis Induction Across Cancer Models

Studies on ferroptosis induction in cultured cells often focus on a handful of lines. Given our failure to induce tumor regression in A549 and HT-1080 cells, we sought to better understand the breadth of responses to ferroptosis inducers across tumor types and in comparison to non-transformed cells. We undertook phenotypic modulatory profiling across 100 cell lines: 70 solid tumor CCLs, 20 myeloid and lymphoid (M&L) CCLs, and 10 non-transformed cell lines (**Figure 3A-B, Supplementary Table 4**). We performed 9-point log_2_ dose-response experiments with RSL-3, erastin, BSO, and titrated exogenous cystine levels; additionally, we co-treated with either the Fer-1 or βMe. For comparison, we included eprenetapopt, a small molecule with GSH-depleting activity^31^ that was evaluated in phase III clinical trials as a treatment for *TP53*-mutant myeloid malignancies^32,33^, but failed to demonstrate sufficient efficacy. We validated these results individually in a subset of 15 cell lines and found highly concordant results (**Extended Data Figure 4A-B, 5A**). We aligned these data with public data reporting drug sensitivity to relevant compounds in these cell lines (CTRPv2^34^) and the dependency on targets GCLC, GPX4, and SLC7A11 (DepMap^35^), and found highly significant concordance, validating the modulatory profiling results (**Extended Data Figure 5B-C**). In general, CCLs are less sensitive to ferroptosis induction compared to non-transformed cell lines, with CCLs derived from solid tumors being more resistant than those from M&L malignancies (**Figure 3C-D**). As previously observed,^36^ eprenetapopt selectively targets M&L CCLs over non-transformed cell lines, supporting the notion our approach can identify such transformation or cell type-specific therapeutic indexes (**Figure 3C**). However, we do not attribute this eprenetapopt selectivity to ferroptosis induction as Fer-1 provides no protection (**Figure 3D**).

**Figure 3.**
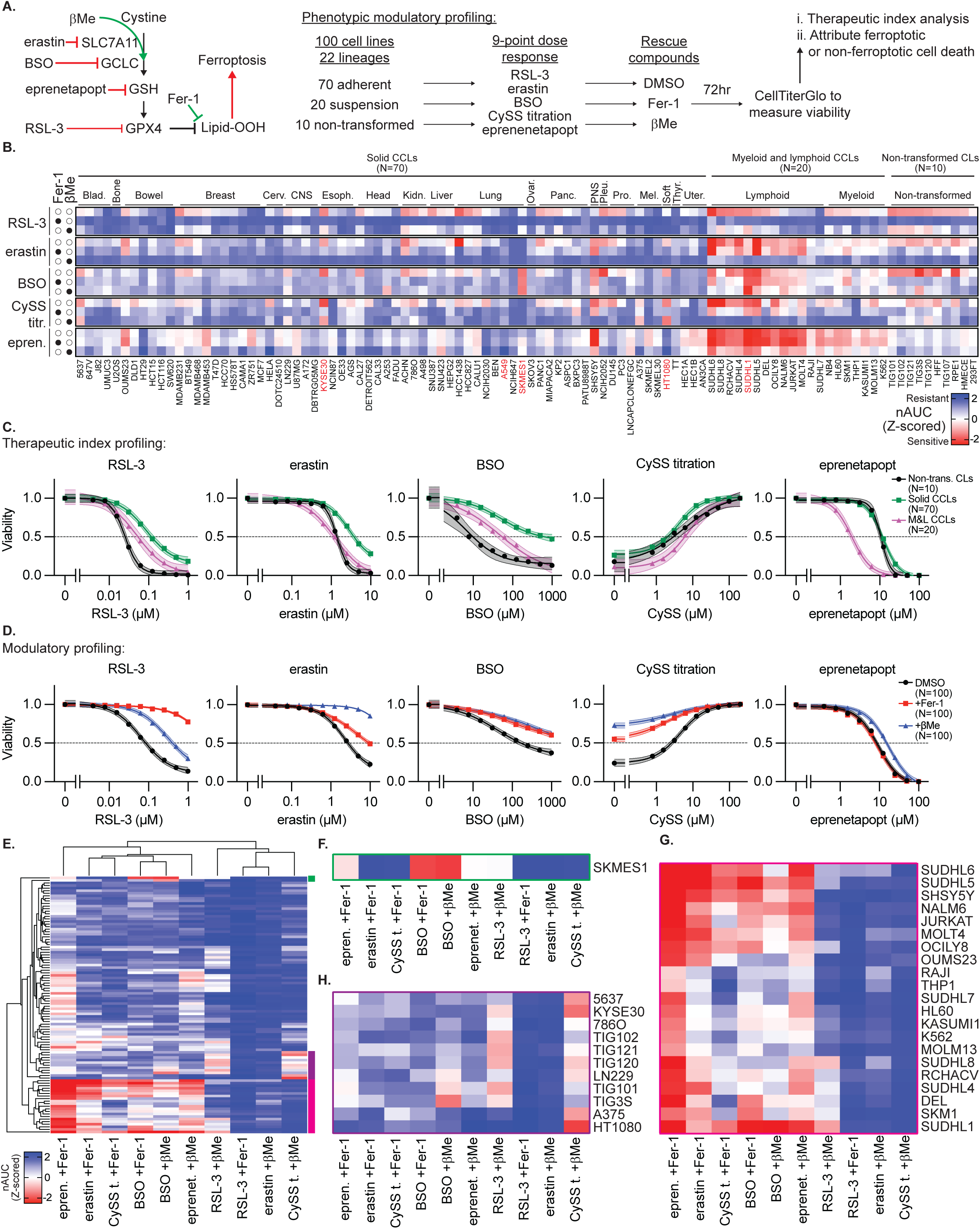
Phenotypic Modulatory Profiling Defines Therapeutic Index of In Vitro Ferroptosis Induction Across Cancer Models. **A**, Schematic outlining phenotypic modulatory profiling approach. 100 cell lines are treated with small molecule inhibitors of SLC7A11, GCLC, GSH, or GPX4, with or without reductant treatment or antioxidant rescue, and viability measurement. **B**, Heatmap reporting Z-score of normalized area under the curve (nAUC) for 9-point dose response of each treatment condition in each cell line. Mean of 2 technical replicates used to generate 1 nAUC biological replicate. Cell lines of subsequent focus in red text. **C-D**, Data from *B* aggregated by indicated categories (*C*: solid tumors, myeloid and lymphoid, or non-transformed cell lines; *D*: DMSO, Fer-1, or βMe treatment). Data report mean ± 95% confidence interval. **E**, Hierarchical clustering of data from *B*. Colored bars at right correspond to detail in *F-H*. **F-H**, Detail of analysis from *E,* indicating cell lines with distinct responses to indicated perturbations.

From these data, we noted a uniformity of responses across approximately two-thirds of cell lines, which exhibit varying degrees of sensitivity to ferroptosis induction that is rescued by Fer-1 co-treatment. Remarkably, we also noticed that βMe can broadly rescue the effects of cystine deprivation and SLC7A11 inhibition (**Figure 3B, 3D**). While prior work has demonstrated that βMe supplementation could bypass SLC7A11-dependent cystine uptake, the presumed mechanism was through reduction of extracellular cystine to cysteine and import through neutral amino acid transporters.^3^ As βMe also prevented cell death in cystine-free conditions, we hypothesized that there exists another mechanism by which βMe and cystine each act to suppress ferroptosis.

Several cell line clusters also exhibited distinct, unexpected response profiles of interest because they were not rescued by Fer-1 and/or βMe (**Figure 3E**). Most notably, one cell line (SKMES1, a small cell lung CCL) exhibits exquisite sensitivity to BSO, even under ferroptosis rescue conditions (**Figure 3F**). Also, a large group of M&L CCLs exhibit sensitivity to BSO treatment, erastin treatment, and cystine withdrawal that is poorly rescued by Fer-1, despite Fer-1 being able to readily rescue the effects of GPX4 inhibition (**Figure 3G**). Finally, HT-1080 cells, frequently used by the ferroptosis field, nucleates a set of cell lines refractory to βMe rescue under cystine deprivation, including several immortalized fibroblast lines and esophageal squamous cell CCL KYSE30 (**Figure 3H**).

These data are consistent with mechanisms operating in responses to canonical ferroptosis inducers that are distinct from ferroptosis. As we struggled to induce tumor regressing by triggering ferroptosis, we explored the anti-cancer impact of inducing non-ferroptotic responses in these models using the corresponding canonical ferroptosis inducers.

## GCLC inhibition in TXNRD1-deficient models drives regulated, non-ferroptotic cell death and causes tumor regression

We confirmed that SKMES1 has high sensitivity to BSO (GI_50_ 2.2 μM) that is not rescued by either Fer-1 or βMe (**Figure 4A**). We also validated that lymphoid cell lines exhibit the most dependence on GSH biosynthetic genes (**Extended Data Figure 6A**), which can be appreciated in an unbiased fashion by the correlation of GCLC dependence with the lymphoid-restricted transcription factor IKZF1 (**Extended Data Figure 6B**), and confirmed with validation by modulatory profiling (**Extended Data Figure 6C**). Of note, these cell lines have universally low SLC7A11 expression (**Extended Data Figure 6D**), concordant with the heighted sensitivity to epnrenetapopt.^37^ SUDHL1, a lymphoid-derived CCL, was chosen as an exemplar of this group (BSO GI_50_ 3.47 μM), and we identified a highly BSO-sensitive mouse lymphoma cell line model (A20 lymphoma cells, BSO GI_50_ 2.22 μM, **Extended Data Figure 6E**), enabling us to assess BSO efficacy in an immunocompetent setting. We engrafted each of these models in immunocompromised mice or syngeneic Balb/c hosts until palpable tumors formed, and randomized tumor-bearing mice to either 50 mM BSO or drinking water. Similar to HT-1080, tumors derived from SUDHL1 and A20 failed to respond to BSO treatment (**Extended Data Figure 6F-G**). Remarkably, BSO was efficacious in controlling the growth of SKMES1, inducing partial or complete regression in 4/5 tumors (**Figure 4B**).

**Figure 4.**
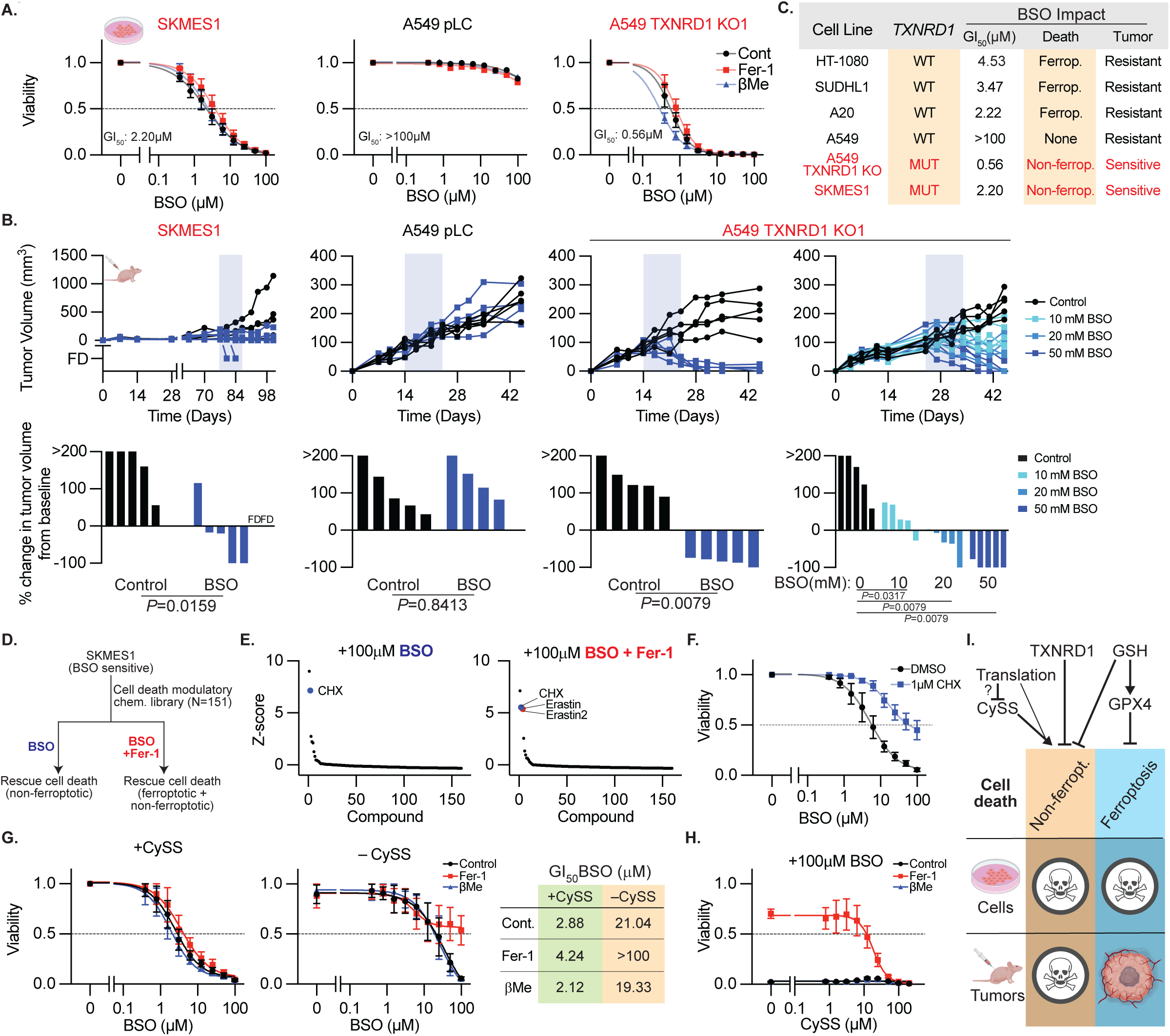
GCLC inhibition in TXNRD1-deficient models drives regulated, non-ferroptotic cell death and causes tumor regression. **A**, Viability (ATP) of SKMES1, A549 cells, or A549 clones in which TXNDR1 is deleted, treated with BSO, with or without βMe (50 μM), or Fer-1 (10 μM), 3 days. Data report mean ± SEM of n=3 biological replicates. GI50 indicated. **B**, Growth of individual subcutaneous tumors derived from cell lines in *A* with or without BSO in drinking water (shaded region) reported as tumor volume (above) or change in tumor volume 21 days after start of BSO treatment or ethical endpoint if reached earlier (below). Data are individual tumor measurements. FD = animal found dead (data censored). P value reports result of Mann-Whitney U test. **C**, Summary of the impact of BSO on models tested. **D**, Schematic of chemical screen for suppressors of non-ferroptotic cell death mechanism. **E**, Z-score of viability (ATP) in SKMES1 cells treated with BSO (100 μM) with or without Fer-1 (10 μM). Cycloheximide (CHX) and erastin or erastin2 indicated (right only). Data report the mean of 2 technical replicates. **F**, Viability (ATP) of SKMES1 cells upon treatment with BSO, with DMSO or CHX (1 μM), 3 days. **G**, Viability (ATP) of SKMES1 cells upon treatment with BSO, with or without CySS supplementation (200 μM), βMe (50 μM), or Fer-1 (10 μM), 3 days. GI50 values from *G* reported in table at right. **H**, Viability (ATP) of SKMES1 cells upon CySS withdrawal and BSO treatment (100 μM), with or without βMe (50 μM), or Fer-1 (10 μM), 3 days. Data from *F-H* report mean ± SEM of n=3 biological replicates. **I**, Model describing the contribution of GSH and thioredoxin to ferroptosis the identified non-canonical cell death pathway, and its regulation by translation and cyst(e)ine availability.

Given the response of SKMES1 tumor xenografts to BSO treatment, we sought to determine what drives the non-ferroptotic cell death observed. Using our phenotypic modulatory screen dataset, we searched public data for biomarkers that predict sensitivity to BSO that is not rescued by Fer-1 or βMe (**Extended Data Figure 7A**).^35^ This analysis revealed that SKMES1 was the only cancer cell line with biallelic inactivating mutations in *TXNRD1*,^35^ which encodes the selenoprotein thioredoxin reductase 1 (TXNRD1), responsible for maintaining the cytoplasmic pool of thioredoxin, a protein-based antioxidant.^38,39^ To verify whether *TXNRD1*-deficiency drives non-ferroptotic cell death in response to BSO, we generated CRISPR-mediated single-cell TXNRD1 KO clones in A549 cells (**Extended Data Figure 7B**). Remarkably, these TXNRD1 KO clones exhibited a 500-fold increase in BSO sensitivity that was not rescued by Fer-1 or βMe, consistent with ferroptosis-independent cell death (**Figure 4A, Extended Data Figure 7C**). Expression of an sgRNA-resistant TXNRD1 cDNA rescues BSO sensitivity in A549 TXNRD1 KO clones and SKMES1 cells (**Extended Data Figure 7D**). However, we did not observe rescue of TXNRD1 deficiency upon expressing TXNRD1 mutants that lack oxidoreductase activity (selenocystine-to-cysteine mutant) or lack the SECIS, confirming that BSO sensitivity depends on the canonical oxidoreductase function of TXNRD1 (**Extended Data Figure 7D**). Expression of a bifunctional glutamate-cysteine ligase/glutathione synthetase enzyme, GshF, from S. thermophilius, which is insensitive to BSO and GSH negative-feedback regulation rescues cell death induced by BSO in TXNRD1 deficient (A549 TXNDR1 KO and SKMES1), and proficient models (HT-1080, **Extended Data Figure 7E-F**)^40^. Similarly, overexpression of GCLC or a BSO-resistant GCLC mutant partially rescues the effect of BSO (**Extended Data Figure 7G**). Concordantly, closely related BSO analogues that fail to deplete GSH (L-buthionine sulfoxide, L-methionine sulfoximide, and L-methionine sulfoxide) do not trigger cell death in in TXNRD1-deficient models or HT-1080 (**Extended Data Figure 7H-K**). Overexpression of GPX4 or the GPX4^U46C^ mutant, which can drive resistance to BSO-induced ferroptosis, does not affect the BSO-sensitivity of TXNRD1-deficient models (**Extended Data Figure 7L**).

Importantly, xenografted tumors derived from TXNRD1 KO A549 cells regress in response to BSO, some completely, and in a dose-dependent manner (**Figure 4B**), consistent with prior work that the synergistic targeting of GCLC and TXNRD1 can limit tumorigenesis.^38,41^ Altogether, these data demonstrate that TXNRD1-deficency drives sensitivity to a GPX4-independent, non-ferroptotic form of cell death. Moreover, triggering this cell death mechanism can regress established tumor xenografts, in contrast to all other BSO-sensitive models tested (**Figure 4C**).

To understand the cell death process triggered by TXNRD1 deficiency and GSH synthesis inhibition, we applied a chemical library of known cell death inhibitors and inducers to SKMES1 cells treated with BSO with or without Fer-1 (**Figure 4D**). The protein synthesis inhibitor cycloheximide rescues the effects of BSO on cell viability in both the presence and absence of Fer-1 (**Figure 4E, Supplementary Table 5**), an effect that we validate in individual experiments (**Figure 4F, Extended Data Figure 8**). In contrast, inhibitors of apoptosis, necroptosis, necrosis, pyroptosis, and ferroptosis all fail to rescue this form of cell death alone or in combination with Fer-1 (**Supplementary Table 5**).

Interestingly, cystine uptake inhibitors erastin and erastin2 also rescue cell viability only in the presence of Fer-1 (**Figure 4E**). Both TXNRD1 and GSH reduce disulfide bonds and convert cystine to cysteine intracellularly^39^, and SKMES1 is the only cell line dependent upon the glutathione reductase, *GSR*.^35^ Therefore, we hypothesized that cystine availability may influence the cytotoxicity of BSO in *TXNRD1*-deficient models. In SKMES1 cells, we compared BSO sensitivity in cystine-replete and depleted conditions, with and without co-treatment with Fer-1 or βMe (**Figure 4G**). While BSO induces non-ferroptotic cell death under cystine replete conditions, cystine withdrawal limits the impact of BSO on cell viability and Fer-1 rescues ferroptosis now induced by combined cystine deprivation and BSO treatment (**Figure 4G**). Remarkably, when we treat cells with a fixed concentration of BSO and titrate cystine out of the media, we observe a dose-dependent rescue of viability in the Fer-1 co-treatment condition (**Figure 4H**). These data demonstrate that cystine depletion suppresses the identified non-ferroptotic cell death mechanism caused by combined TXNRD1/GCLC deficiency. This result is particularly surprising in that cystine withdrawal cannot restore GSH levels, despite intracellular GSH production suppressing this cell death mechanism (**Extended Data Figure 7F**). Together, these data demonstrate that combined deficiency in the cytosolic thioredoxin reductase system and pharmacologic GCLC inhibition triggers a form of cell death that is distinct from ferroptosis, regulated by cystine availability and protein synthesis, and can be triggered both in cultured cells and in tumor xenografts (**Figure 4I**).

## Maintaining Selenoprotein and GPX4 Function Bypasses the Requirement for Exogenous Cystine

As described above, βMe was surprisingly capable of broadly inhibiting cell death induced by cystine depletion, including in the vast majority of solid, M&L CCLs and non-transformed CLs (**Figure 5A-B**). Therefore, we generated SLC7A11 KO clones in A549 cells, which require βMe to sustain viability and suppress ATF4-mediated stress responses (**Figure 5C, Extended Data Figure 9A**), consistent with βMe driving cystine to cysteine conversion and SLC7A11-independent transport. Interestingly, SLC7A11 KO clones require nanomolar concentrations of βMe to sustain viability, while protection from cystine deprivation requires micromolar βMe concentrations, implying a distinct mechanism (**Extended Figure 9B**). Moreover, while thiol-containing dithiothreitol (DTT) or tris(2-carboxyethyl) phosphine (TCEP) also bypass the requirement for SLC7A11, they cannot protect against cystine deprivation (**Extended Data Figure 9C**). βMe supplementation fails to restore GSH levels, which are well-suppressed, disconnecting GSH level from proliferative capacity (**Extended Figure 9D**). While mammalian cells have the capacity to generate cysteine *de novo* through transsulfuration of serine,^42^ tracing of uniform isotopically labelled serine (U-^13^C-Serine) failed to detect transsulfuration pathway activity (**Extended Data Figure 9E**), in agreement with the observed loss of GSH (**Extended Figure 9D**), as well as previous reports indicating a lack of transsulfuration pathway activity in cultured cancer cells.^43^ The ability of βMe to rescue cell viability in cystine-free conditions despite GSH depletion is consistent with our observations that βMe restores exponential growth of cells lacking GCLC (**Figure 2D,G**) and broadly protects BSO-treated cells from ferroptosis (**Figure 3D**). Remarkably, 50 μM βMe supplementation supports long-term culturing of SLC7A11 KO clones both in the presence and absence of extracellular cystine, and the long-term culturing of WT cells in cystine-free media (**Figure 5D**). These data indicate that βMe can promote SLC7A11-independent cyst(e)ine uptake and cystine-independent growth through different mechanisms and demonstrate that βMe can replace extracellular cystine *in vitro* to promote robust cell proliferation.

**Figure 5.**
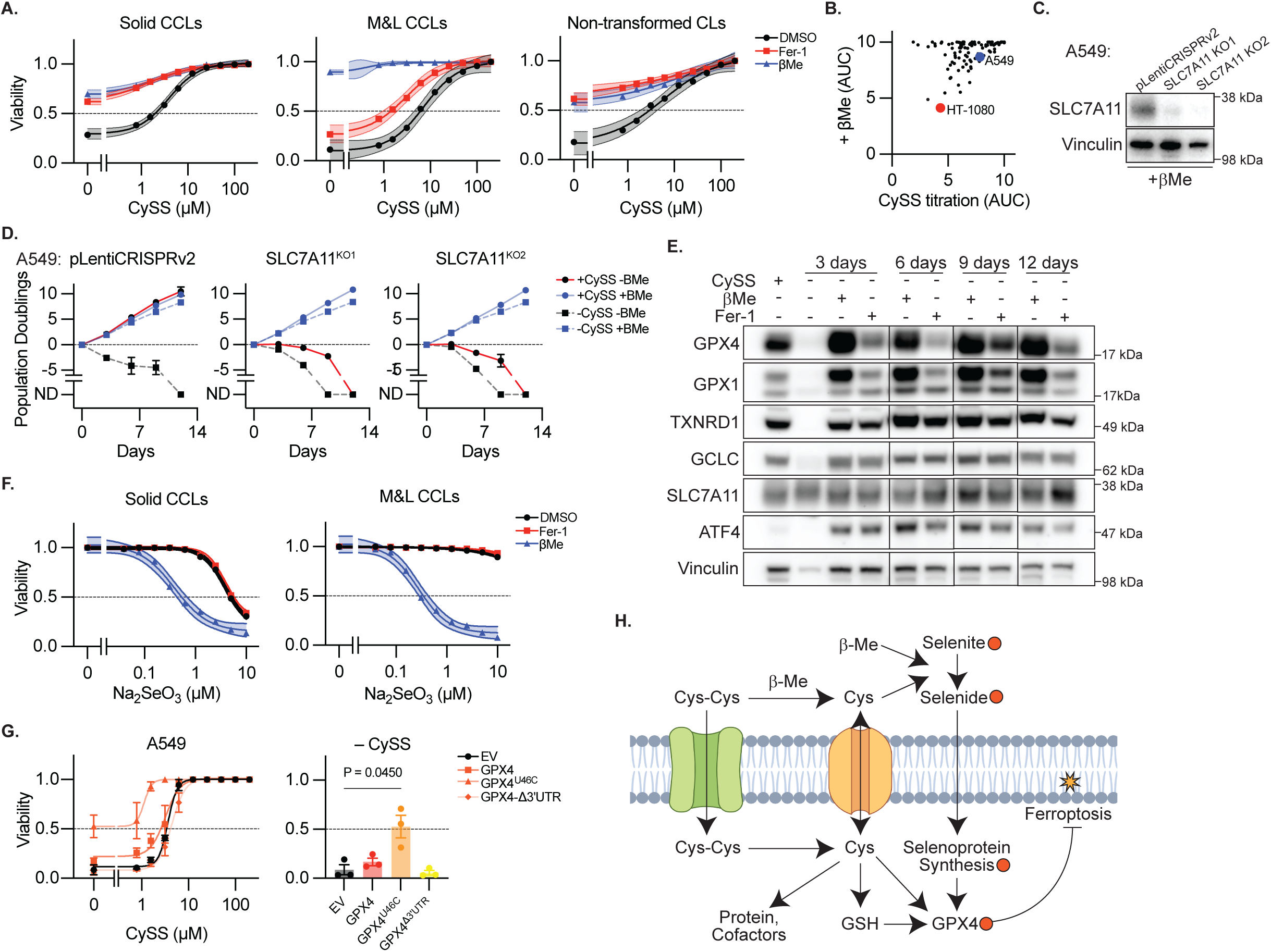
Maintaining Selenoprotein and GPX4 Function Bypasses the Requirement for Exogenous Cystine. **A**, Sensitivity to CySS deprivation from Figure 3B aggregated by indicated categories (solid, myeloid and lymphoid, or non-transformed cell lines). Data report mean ± 95% confidence interval. **B**, Cell line sensitivity to CySS deprivation and impact of βMe addition (AUC). A549 and HT-1080 cell lines indicated. **C**, Immunoblot for indicated proteins from lysates of A549 cells and two clones in which SLC7A11 is deleted, maintained in βMe. Inferred MW of band indicated. **D**, Cumulative population doublings of cell lines from *C* cultured in the presence or absence of CySS and βMe. ND, no remaining detectable cells. Data report mean ± SEM of n=3 biological replicates. **E**, Immunoblot for indicated proteins from lysates of A549 cells cultured in the presence of absence of CySS (200 μM), βMe (50 μM), or Fer-1 (10 μM). Inferred MW of band indicated. **F**, Viability (ATP) aggregated by indicated category of cell lines treated with Na2SeO3 and DMSO, βMe (50 μM), or Fer-1 (10 μM), 3 days. Data report mean ± 95% confidence interval of n=10 solid and n=5 M&L cell lines (from Extended Figure 4), each in technical duplicate. **G**, Viability (ATP) of A549 cells expressing the indicated GPX4 cDNAs and selenocysteine (U46C) or SEICS (Δ3’UTR) mutants, cultured in indicated CySS conditions, 3 days. Bar graph reports data from line graph at 0 μM CySS. Data report mean ± SEM of n=3 biological replicates. P value reports result of heteroscedastic student’s t-test. **H**, Model describing the role of cyst(e)ine and βMe in supporting selenium import to enable GPX4 function.

In the pursuit of other mechanisms by which βMe could prevent ferroptosis, we turned our focus to selenium import, as previous reports have suggested that SLC7A11 function and cystine uptake maintains an extracellular redox environment that favors selenium uptake^44,45^. Selenium is an essential nutrient for selenocysteine synthesis that is either added to media in the form of sodium selenite, or otherwise present in serum selenoproteins or other small molecular weight selenium species. Selenite reduction to selenide by extracellular thiols promotes its import and can drive selenium toxicity.^44^ Consistent with this mechanism, there are only modest cell viability effects observed for SLC7A11 loss in pooled loss-of-function screening datasets (e.g. DepMap). Moreover, co-culture transwell assays revealed that control cells, but not SLC7A11 KO cells, can support the proliferation of SLC7A11 KO cells, demonstrating that a SLC7A11-dependent component of the conditioned media can overcome the SLC7A11 requirement (**Extended Data Figure 9F**). Consistent with the hypothesis that cystine deprivation limits selenium availability, we find that cystine depletion results in loss of selenoproteins sensitive to selenium withdrawal (GPX4 and GPX1), which can be robustly rescued by βMe supplementation, even after 12 days of exponential proliferation (**Figure 5E**), and upon SLC7A11 KO (**Extended Data Figure 9G**). βMe supplementation increases the cytotoxic effects of sodium selenide treatment across multiple cell line models (**Figure 5F**) to a much greater extent than DTT and TCEP (**Extended Data Figure 9H**), consistent with βMe promoting selenium uptake. Importantly, overexpression of a selenium-independent GPX4 mutant (GPX4^U46C^) maintains cell viability under cystine-deprivation, while wild-type GPX4 does not (**Figure 5G**).

Further supporting the hypothesis that cystine supplementation enables selenium availability for GPX4 function, in the small number of models where βMe does not protect against cystine deprivation (e.g. HT-1080 and KYSE30, **Figure 3H, 5B**), βMe can still maintain selenoprotein levels, yet GPX4^U46C^ overexpression fails to prevent ferroptosis (**Extended Data Figure 10A-B**). These data are consistent with a heightened requirement for cystine or GSH levels to block ferroptosis, or the heightened requirement for selenoproteins other than GPX4 in these models. CRISPR-based screening and validation experiments emphasize that inhibition of PUFA incorporation into membrane lipids via suppression of LPCAT3 readily blocks ferroptosis induced by SLC7A11 KO or cystine deprivation in A549 cells, while minimally affecting HT-1080 and KYSE30 cell lines (**Extended Data Figure 10C-D, Supplementary Table 6**), further demonstrating the requirement for cystine beyond supporting GPX4 function in this small subset of models.

These data demonstrate that the primary function of exogenous cystine in supporting the viability of cultured cells is via supporting selenium import and selenoprotein function, as cells can survive long-term culture in the absence of cystine and maintain proliferative capacity if selenoprotein function is restored (**Figure 5H**).

## Discussion

Successful development of targeted anti-cancer therapies in the age of precision medicine relies on appropriate target identification and validation, performed in pre-clinical models of cancer. The primary modes of failure of a novel anti-cancer therapy’s path toward clinical approval are often due to limited efficacy and intolerable toxicity.^46^ The financial costs of drug development, together with the enormous risks and burdens facing patients entering clinical trials, underscores the critical importance of diligent pre-clinical target validation before promoting a new cancer treatment modality into clinical investigation.

Over two decades has passed since the first ferroptosis inducers were identified, and while extraordinary gains have been made in defining and delineating the *in vitro* molecular mechanisms of ferroptosis as a cell death process, less progress has been made in translating these insights into clinically useful ferroptosis inducers. Poor solubility and pharmacokinetic profiles have limited the direct application of erastin and RSL-3; however, efforts to generate bioavailable SLC7A11 and GPX4 inhibitors have succeeded in target engagement but failed to induce tumor control without on-target off-tumor toxicity.^47^ For example, Abbvie scientists developed compound 24, an RSL-3 analogue with appropriate pharmacokinetics and demonstrated *in vivo* target engagement.^48^ However, this compound resulted in on-target kidney toxicity and an absence of anti-tumor efficacy in animal models.^48^ These results were in keeping with genetic ablation data in mice which reported kidney toxicity associated with whole body, inducible GPX4 depletion.^49^ Additionally, GCLC inhibitor BSO has been used in humans, and has been tested extensively in pre-clinical tumors models in combination with chemotherapy.^50^ BSO was shown to induce modest tumor growth inhibition as a single agent by Kojin Therapeutics, an effect attributed to ferroptosis induction.^51^ However, clinical testing has resulted in two deaths in pediatric patients without substantial anticancer activity.^52^ These results indicate that on-target renal toxicity will limit cancer therapy via single-agent inhibition of the SLC7A11-GSH-GPX4 axis.

Our results highlight that cancer models capable of undergoing ferroptosis robustly in cell culture fail to modulate tumor growth in mice. Moreover, several unbiased *in vivo* CRISPR screens utilizing metabolic libraries which include ferroptosis targets, *Gpx4*, *Gclc, Slc7a11*, and *Aifm2* report no significant changes to guide abundance.^53,54^ Recent evidence demonstrates that 2D culture itself enhances the PUFA composition of membrane lipids, artificially enhancing sensitivity to GPX4 inhibition.^55^ Furthermore, the cystine:cysteine redox couple is far more oxidized in standard cell culture media than in vivo, including in tissues and tumors.^56^ These findings come against the backdrop of a clear pro-tumor effect of antioxidant supplementation in both mice and humans,^57,58^ as well as the clear activation of the NRF2 antioxidant response pathway through recurrent mutations of KEAP1, NFE2L2 (encoding NRF2) and CUL3 in lung adenocarcinoma, lung squamous cell carcinoma, head and neck squamous cell carcinoma, esophageal squamous cell carcinoma, and others, while non-genetic NRF2 pathway activation is observed in additional tumor types.^59^ NRF2 activates several genes that protect against ferroptosis, including those encoding the iron storage protein ferritin, the GSH biosynthesis pathway, and those promoting NADPH production or recycling.^59^ However, while NRF2 activation clearly drives tumorigenesis in mice and humans, it is less certain whether sustained NRF2 pathway activation is important for tumor maintenance.

These considerations lead to several points that might explain the diversity of results from different laboratories studying ferroptosis in tumor models. First, there may be a differential requirement for ferroptosis protection during tumor initiation, progression, and metastasis. Indeed, many genetically-driven preclinical models test target validation by suppressing the target prior to or concomitant with tumor engraftment, as opposed to inhibiting the target in an inducible fashion in established tumors, which would better model potential future drug treatment. Groups have also identified metastatic environments where iron levels are limited, including lymphatic vessels, which impact oxidative stress sensitivity.^60^ Second, the tumor types in which NRF2 pathway alterations arise may indicate the importance of anatomic location in driving oxidative stress sensitivity. From a very simple perspective, the respiratory airways have increased exposure to environmental O_2_, and these environments can be more hostile to cancer cells that exhibit O_2_ sensitivity.^28^ Neurons may also have heightened sensitivity to ferroptosis as mutations in GPX4 have been identified in humans with the neurodegenerative disease Sedaghatian-type spondylometaphyseal dysplasia and increased ferroptotic-like cell death has been reported in cell and animal models of this disease.^61^ Third, determining whether reduced tumor growth reflects ongoing cell death via ferroptosis is extremely difficult, and for simplicity the field often conflates an increase in tumor growth upon antioxidant treatment with evidence of ferroptosis.^62^ Indeed, there are certainly non-ferroptotic reasons for which antioxidant treatment may rescue the effects of target gene suppression and enhance tumorigenesis, even for molecules classified as ferrostatins. Indeed, recent evidence indicates that some of these molecules act via iron chelation.^63^ Efforts to engineer models that are unable to undergo ferroptosis via inhibition of PUFA incorporation into plasma membranes may be useful in disentangling these phenomena, as was recently shown for an assessment of the ability of FSP1 inhibition to induce ferroptosis in lung cancer.^64^

Given the prominent role of cystine import in our understanding of ferroptosis, our results provide important context for the role of cystine and GSH in cultured cells. We find that cystine can be removed completely from cell culture media in many contexts if cells are provided with another thiol reductant, βMe, permitting long-term exponential growth. These data indicate that imported cystine is exported from cells as cysteine or GSH, where it conditions the media to permit selenium uptake, as indicated by the work of Kim and colleagues.^65^ Indeed, we find that expression of a selenium-independent GPX4 cDNA can support cell viability in the absence of exogenous cystine. The ability of cells to survive long-term culture in cystine free medium is surprising and indicates that cultured cells can obtain sufficient cysteine to support protein synthesis and minimal levels of GSH production from another source, most likely catabolism of extracellular protein. Moreover, our data demonstrate that exponential cell growth does not require the millimolar concentrations of GSH present in cells, as βMe or Fer-1 permit exponential growth of GCLC knockout cells. These results are concordant with the failure of SLC7A11 and GCLC to score as essential in pooled genetic screens.^35^ Indeed, we find that SLC7A11 expressing cells or their conditioned media can readily rescue SLC7A11 knockout cell growth. Thus, we conclude that the principal cell-essential function of cystine in cell culture media (and therefore of SLC7A11 expression in cultured cells) is to support the production of extracellular thiols that permit selenium uptake for the function of selenoproteins like GPX4, in nearly all models.

Our work also identifies rare models with a heightened sensitivity to GCLC inhibitor BSO. While in some models this sensitivity is due to a heightened requirement for GSH in suppressing ferroptosis or supporting selenoprotein function, we find an additional model in which TXNRD1 deficiency drives heightened BSO sensitivity. Interestingly, loss of the cytosolic thioredoxin reductase combined with pharmacologic GCLC inhibition results in a form of non-ferroptotic cell death that can be regulated by extracellular cystine deprivation and protein synthesis inhibition. The rescue of this cell death mechanism by cystine withdrawal is particularly surprising given that BSO triggers GSH depletion and forced GSH restoration rescues this cell death, but cystine withdrawal cannot restore GSH levels, yet is cytoprotective. We speculate that loss of both TXN and GSH systems limits the ability of the cell to buffer intracellular redox states, rendering cells sensitive to extremes in thiol concentrations.

As this non-ferroptotic form of cell death induced by BSO in the context of TXNRD1 loss can readily regress established tumors, our results suggest TXNRD1-deficiency could provide a tumor-intrinsic therapeutic index for the use of agents targeting GCLC, in agreement with previous studies.^38,41^ However, *TXNRD1* mutations emerge rarely in human cancers based on cell line and human tumor biobank analysis, while combined inhibition of both pathways would likely result in on-target toxicity to achieve tumor responses.

Our data demonstrate that the *in vitro* effects of ferroptosis induction strongly over-estimate the impact on tumor control. Therefore, our study suggests that caution should be taken when considering the attribution of ferroptosis induction in cancer cells as a driver of tumor control and regression. Preclinical models of cancer wherein the suppression of the SLC7A11-GCLC-GPX4 axis in cancer cells are invoked as the primary mode of tumor control should be examined to reconsider whether appropriate emphasis is placed on the contribution of ferroptosis to any observed anti-tumor effects.

---

**Extended Data Figure 1.**
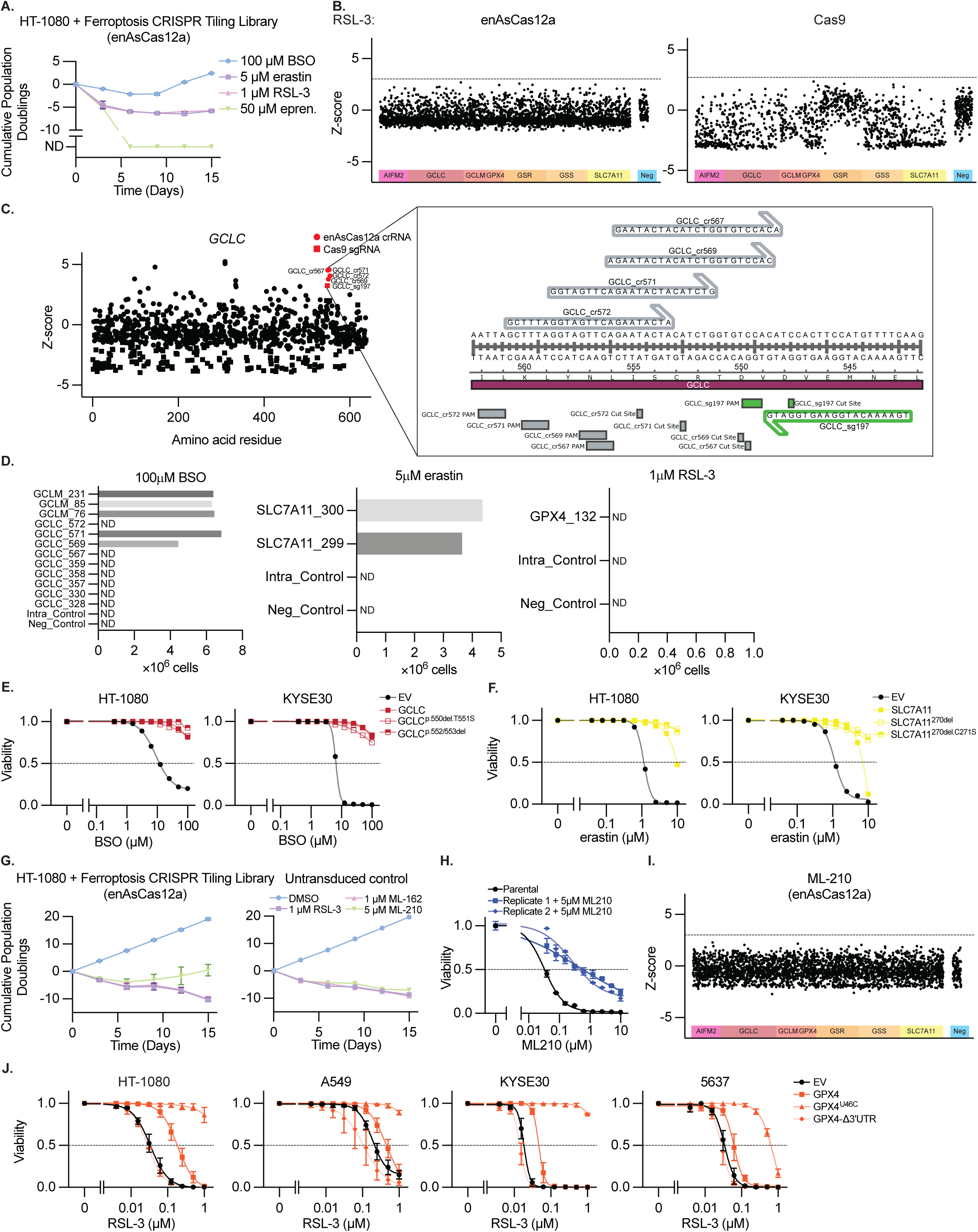
Data Supporting Figure 1 **A**, Cumulative population doublings of HT-1080 cells transduced with the CRIPSR tiling library described in Figure 1 upon selection with BSO (100 μM), erastin (5 μM), RSL-3 (1 μM) or eprenetapopt (50 μM). ND, no remaining detectable cells. Data report mean ± range of n=2 biological replicates. **B**, Z-score of guide RNA enrichment using enAsCas12a (left) or Cas9 (right), tiled across genes indicated upon RSL-3 (1 μM) selection, n=2 biological replicates. **C**, Detail of data from Figure 1B-C showing guide RNAs targeting GCLC enriched upon BSO treatment. Inset shows nucleotide-level resolution of a cluster of highly selected guide RNAs at the GCLC:GCLM binding interface. **D**, Validation experiments in which HT-1080 cells are transduced with individual highly enriched guide RNAs from the tiling screen described in Figure 1 B-C. Data reports number of HT-1080 cells remaining after 15 days treatment with the indicated concentration of BSO, erastin, or RSL-3, 3 days growth, ND = no remaining detectable cells. Data report a single experiment. **E-F**, Viability (ATP) of HT-1080 or KYSE30 cells expressing GCLC, SLC7A11, or cDNAs that mutate regions targeted by positively selected guide RNAs from Figure 1F or 1I, upon treatment with BSO (*E*) or erastin (*F*), 3 days. Data are mean ± SEM, n=3 biological replicates. **G**, Cumulative population doublings of HT-1080 cells transduced with the CRIPSR tiling library described in Figure 1 or parent cell line upon selection with RSL-3 (1 μM), ML-162 (1 μM), or ML-210 (5 μM). Data report mean ± SEM of n=2 biological replicates. **H**, Viability (ATP) of HT-1080 cells selected with ML-210 (*G*) upon treatment with ML-210, 3 days. Data are mean of two technical replicates ± SD. **I**, Z-score of guide RNA enrichment using enAsCas12a, tiled across genes indicated upon ML-210 (5 μM) selection, n=2 biological replicates. **J**, Viability (ATP) of indicated cell lines expressing indicated GPX4 cDNAs and selenocysteine (U46C) or SEICS (Δ3’UTR) mutants, upon RSL-3 treatment, 3 days. Data report mean ± SEM of n=3 biological replicates.

**Extended Data Figure 2.**
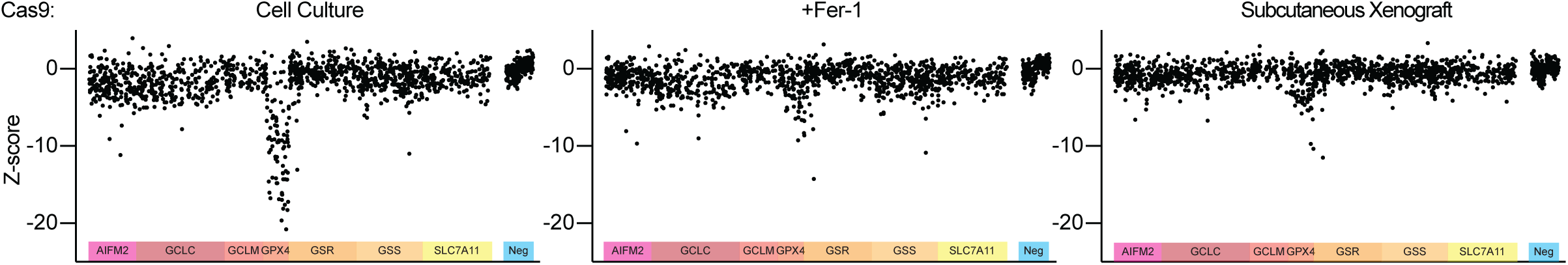
Data Supporting Figure 2A-B. Z-score of guide RNA enrichment using Cas9, tiled across genes indicated upon 15 days *in vitro* culture, including Fer-1 (10 μM) treatment, or upon growth as subcutaneous tumor xenografts (14 days), HT-1080 cells. Data are the mean of individual sgRNAs from 2 biological replicates.

**Extended Data Figure 3.**
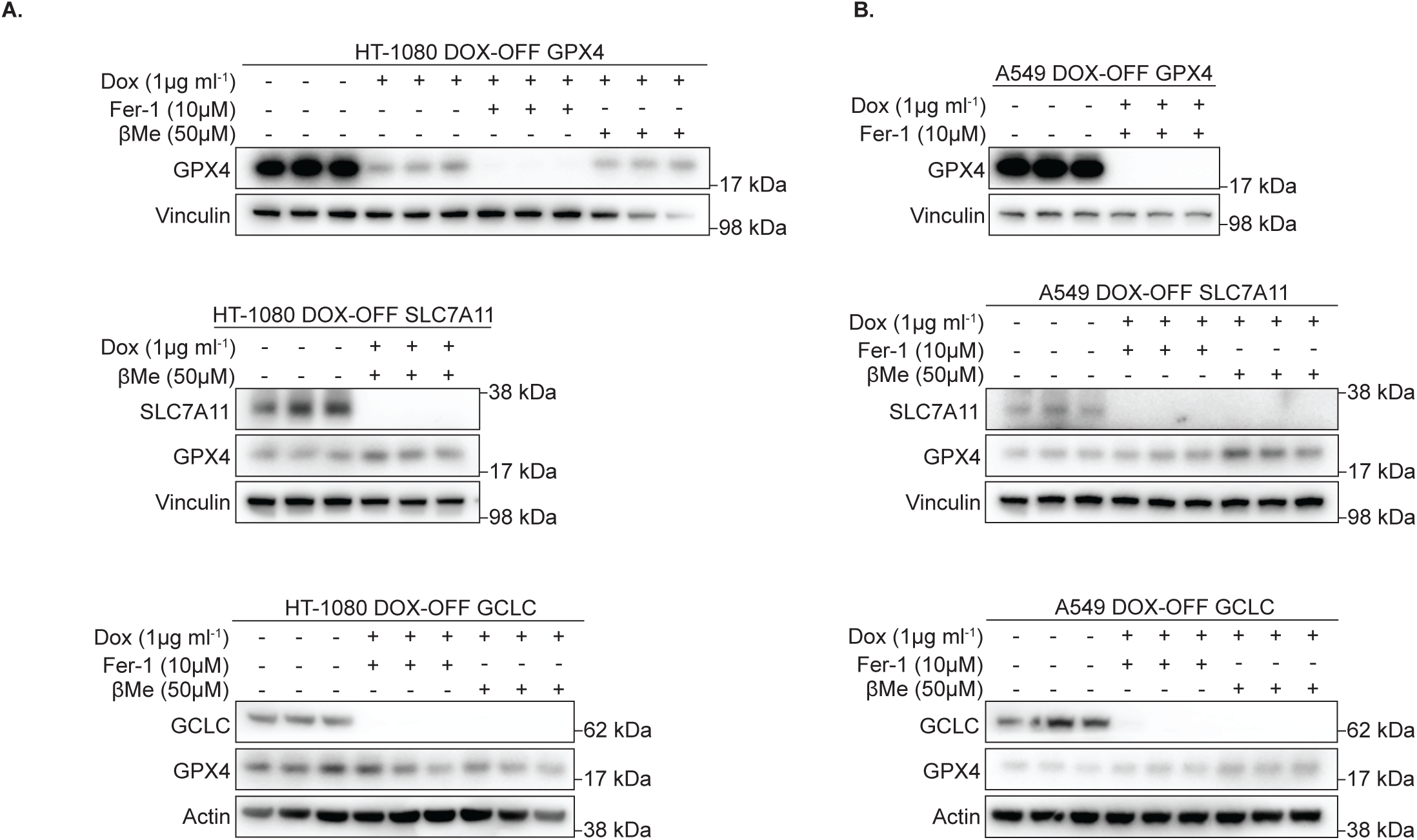
Validation of doxycycline-repressible cell line models. **A-B**, Immunoblots from lysates of HT-1080 (*A*) or A549 (*B*) clones expressing doxycycline (DOX)-repressible cDNAs encoding indicated genes [GPX4 (top), SLC7A11 (middle), or GCLC (bottom)] with CRISPR/Cas9-mediated deletion of the endogenous gene, cultured in the indicated concentrations of DOX, ferrostatin (Fer-1) or βMe. MW of marker indicated.

**Extended Data Figure 4.**
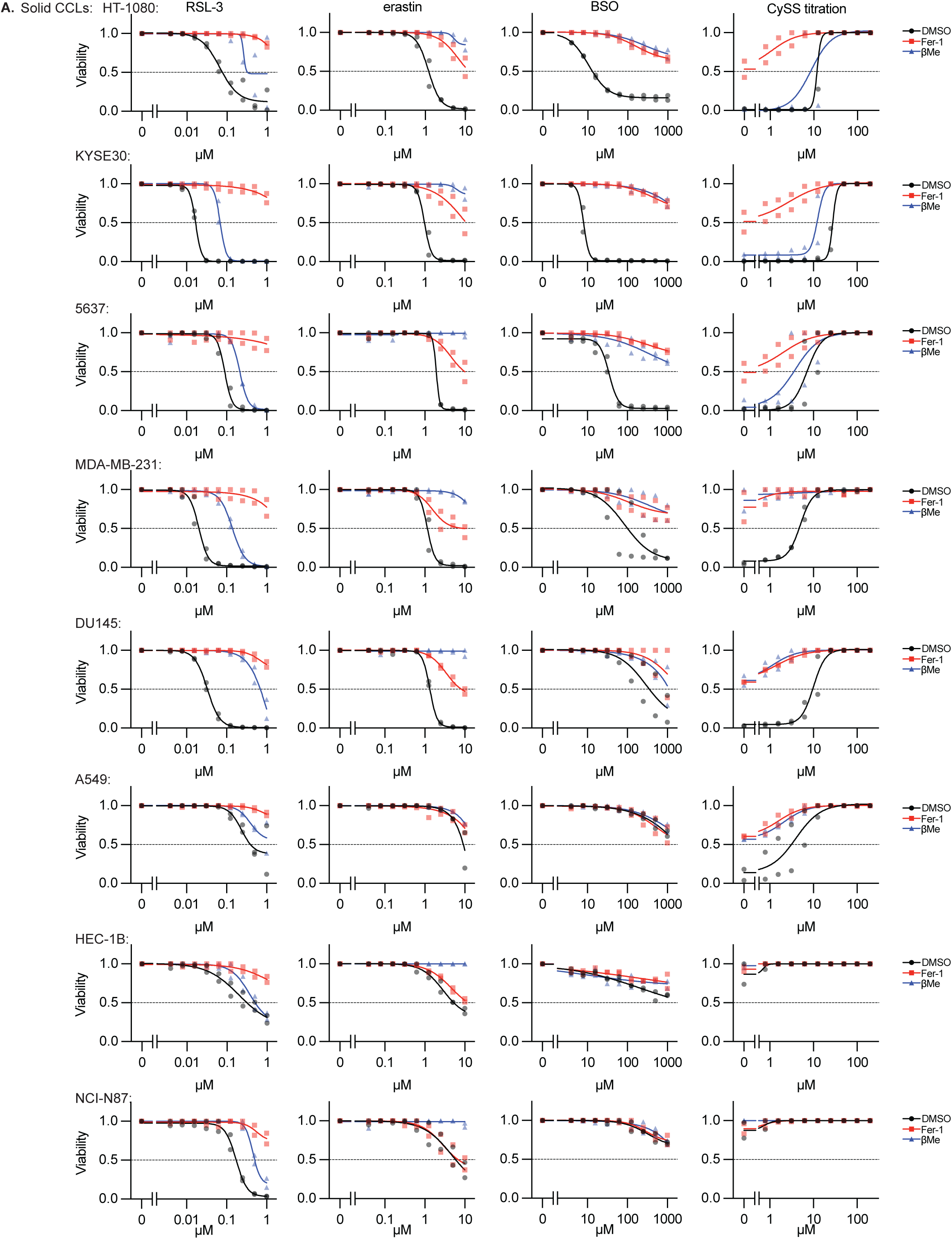
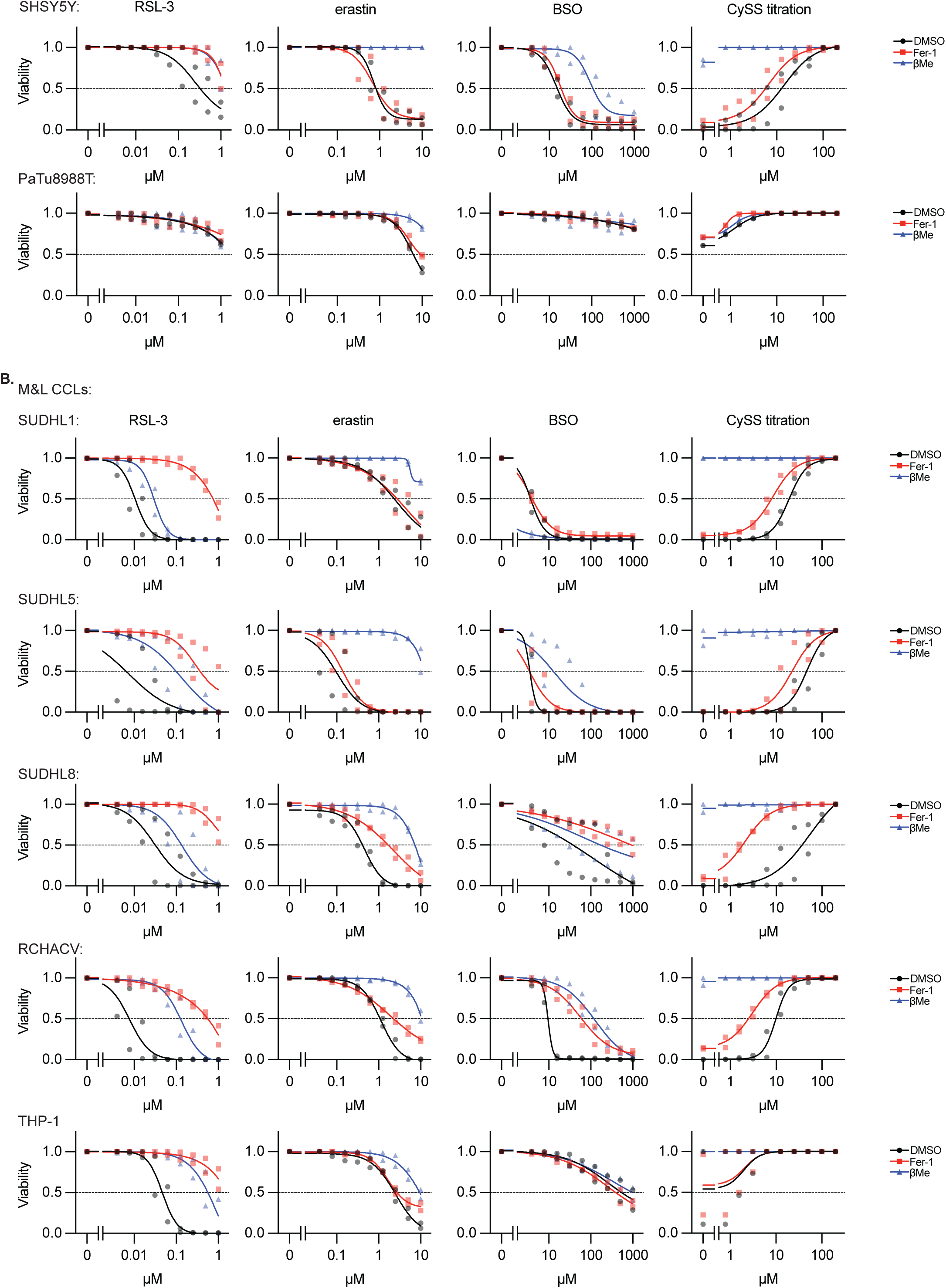
Representative dose responses across 15 cancer cell lines from cell line screen and validation. **A-B**, Viability (ATP) of indicated solid tumor (*A*) or myeloid and lymphoid (*B*) cell lines treated with indicated compounds or in the indicated conditions (RSL-3, erastin, BSO, CySS withdrawal), plus co-treatment with DMSO, βMe (50 μM), or Fer-1 (10 μM), 3 days. Data report mean from two technical replicates from individual experiments for two biological replicates.

**Extended Data Figure 5.**
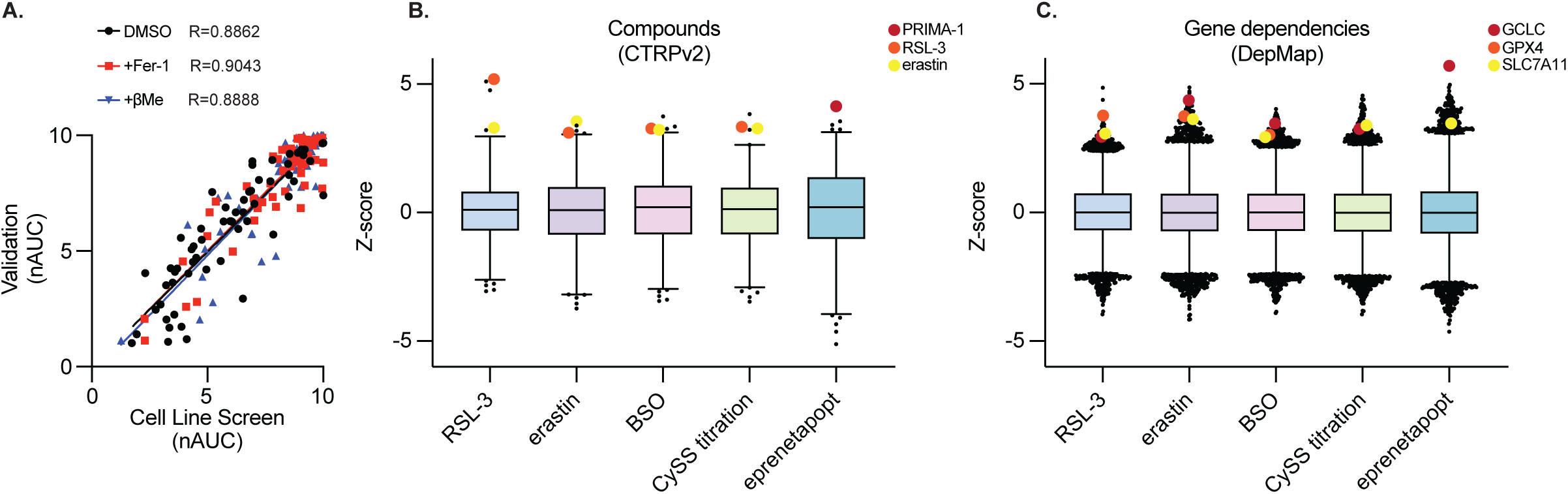
Data Supporting Cell Line Modulatory Profiling. **A**, Comparison of nAUC measurements from Figure 3B and validation experiments in 15 cell lines (Extended Data Figure 4, Mean of 4 technical replicates used to generate 2 nAUC biological replicates). **B-C**, Comparison of data from Figure 3B with (*B*) sensitivity to compounds from Cancer Therapeutics Response Portal or (*C*) gene dependency score (DepMap), relevant compounds and genes indicated (PRIMA-1: eprenetapopt analogue).

**Extended Data Figure 6.**
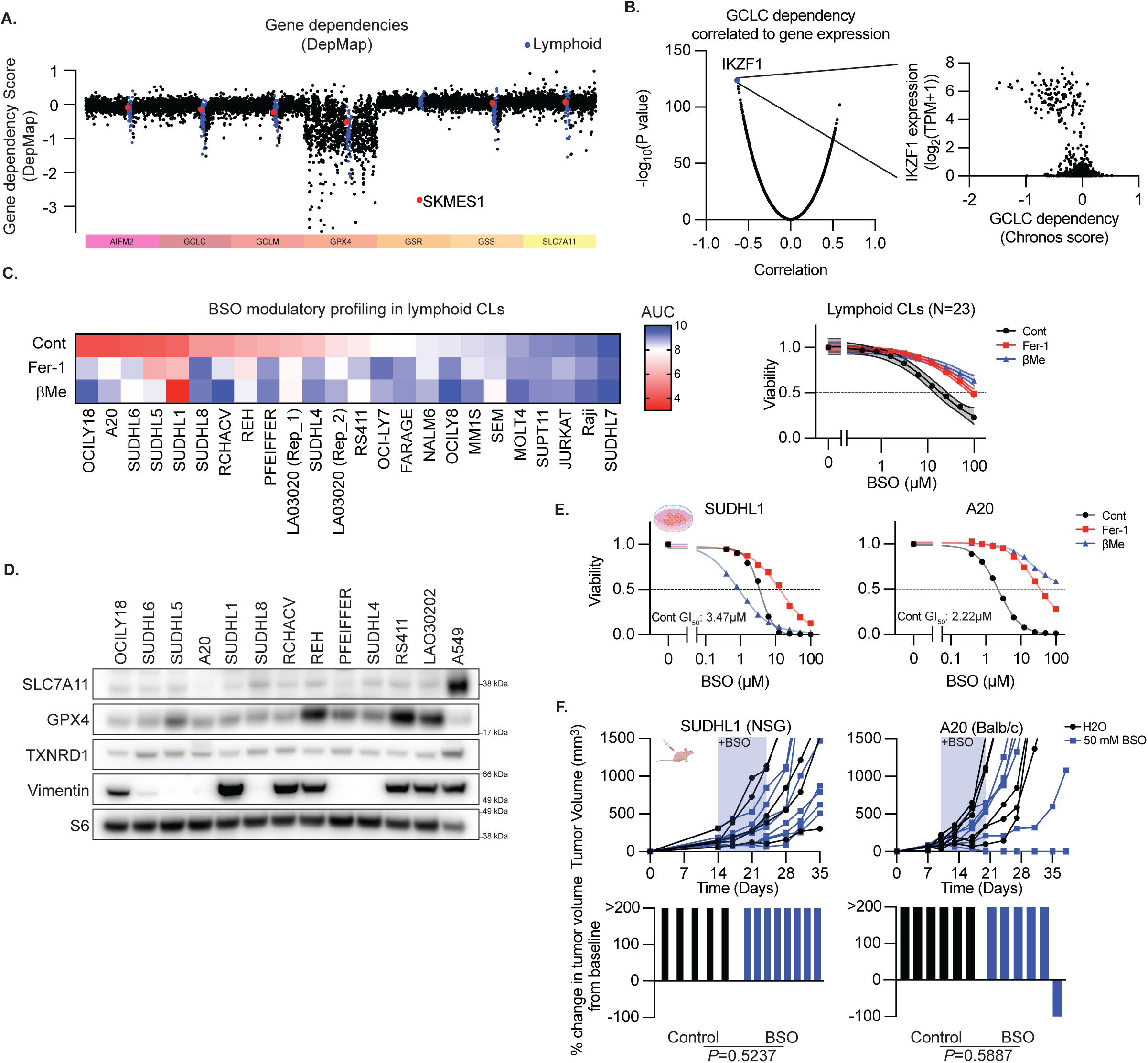
Lymphoid cancer cell lines exhibit *in vitro*-specific BSO sensitivity. **A**, Gene dependency score for the genes indicated (colored bar at bottom) in cell lines from the cancer cell line encyclopedia (DepMap), ordered by tumor type. SKMES1 (red) and lymphoid cell lines (blue) highlighted. **B**, Correlation and significance score of dependency to GCLC loss from the DepMap dataset, highlighting the correlation of IKZF1 to GCLC sensitivity. Inset shows the correlation between IKZF1 expression and GCLC dependency from this dataset. **C**, Left, Heatmap reporting area under the curve (AUC) for 9-point dose response to BSO treatment. Right, aggregated by indicated categories (DMSO, Fer-1, or βMe treatment). Data report mean ± 95% confidence interval. **D**, Immunoblots from lysates of indicated cells lines for indicated proteins. MW of marker indicated. **E**, Viability (ATP) of SUDHL1 or A20 cells treated with BSO, with or without βMe (50 μM), or Fer-1 (10 μM), 3 days. Data report mean of two technical replicates. GI50 indicated **F**, Growth (of individual subcutaneous tumors derived from SUDHL1 or A20 cells with or without BSO in drinking water (50 mM, shaded region) reported as tumor volume (top) or change in tumor volume 21 days after start of BSO treatment or ethical endpoint if reached earlier (bottom). Data are individual tumor measurements. SUDHL1 control, n=5; SUDHL1+BSO n=8; A20 n=6 each group. P value reports result of Mann-Whitney U test.

**Extended Data Figure 7.**
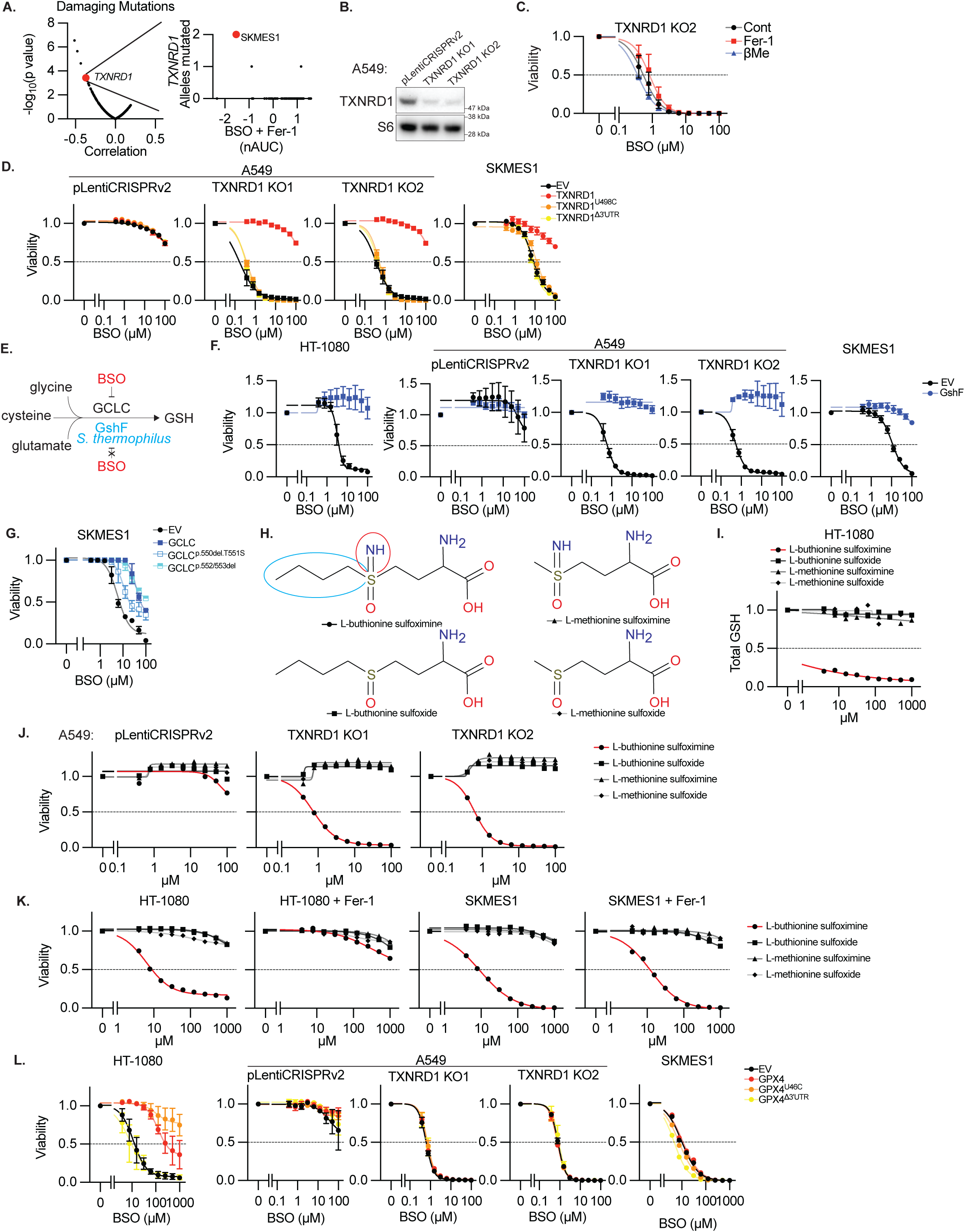
Functional TXNRD1 deficiency renders cells sensitive to BSO. **A**, Left, correlation between the reported number of damaging mutations in each gene (DepMap) and nAUC measurement for BSO sensitivity (+Fer-1 condition, Figure 3B) and associated P value, *TXNRD1* highlighted. Right, correlation between number of number of damaging mutations in *TXNRD1* in each cell line and corresponding nAUC for BSO sensitivity (+Fer-1), SKMES1 cell line highlighted. **B**, Immunoblot for indicated proteins from lysates of A549 cells and two clones in which TXNDR1 is deleted. MW markers indicated. **C**, Viability (ATP) of a TXNRD1 KO clones from *B* upon treatment with BSO, with or without βMe (50 μM), or Fer-1 (10 μM), 3 days. Data report mean ± SEM of n=3 biological replicates. **D**, Viability (ATP) of A549 cells, A549 TXNRD1 knockout clones, or SKMES1 cells expressing the indicated TXNRD1 cDNAs and selenocysteine (U498C) or SEICS (Δ3’UTR) mutants, treated with BSO as indicated, 3 days. Data report mean ± SEM of n=3 biological replicates. **E**, Schematic of glutathione synthesis and BSO sensitivity for GCLC versus GshF. **F**, Viability (ATP) of HT-1080, A549, A549 TXNRD1 knockout clones, or SKMES1 cells expressing S. thermophilus GshF, treated with BSO as indicated, 3 days. Data report mean ± SEM of n=3 biological replicates. **G**, Viability (ATP) of SKMES1 cells expressing exogenous GCLC cDNA and GCLC mutants resistant to BSO identified in Figure 1, treated with BSO as indicated, 3 days. Data report mean ± SEM of n=3 biological replicates. **H**, Schematic of BSO (L-buthionine sulfoximine) and sulfoxide or methionine derivatives. Differences in buthionine vs methionine (blue circle) or sulfoximine vs sulfoxide (red circle) indicated. **I**, GSH content of HT-1080 cells upon treatment with BSO (red line) and indicated related species, 3 days. Data are mean of two technical replicates. **J-K**, Viability (ATP) upon treatment with BSO (red line) and indicated related species for A549 or A549 TXNRD1 knockout clones (*J*), or HT-1080 and SKMES1 cells, including additional Fer-1 treatment (10 μM, *K*), 3 days. Data are mean of two technical replicates. **L**, Viability (ATP) of HT-1080, A549, A549 TXNRD1 knockout clones, or SKMES1 cells expressing the indicated GPX4 cDNAs and selenocysteine (U46C) or SEICS (Δ3’UTR) mutants, treated with BSO as indicated, 3 days. Data report mean ± SEM of n=3 biological replicates.

**Extended Data Figure 8.**
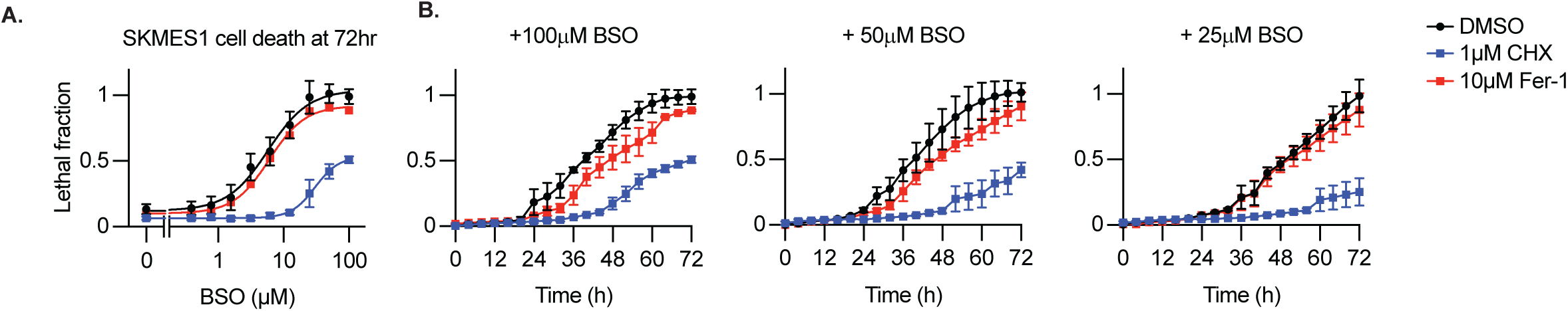
Kinetics of Cell Death in SKMES1 Cells. **A**, Lethal fraction (propidium iodine uptake) of SKMES1 cells treated with indicated concentrations of BSO with or without vehicle (DMSO), cycloheximide (1µM) or Fer-1 (10µM) co-treatment, 72hrs. **B**, Kinetics of cell death in SKMES1 cells treated at 0hrs with indicated concentrations of BSO with or without vehicle (DMSO), cycloheximide (1µM) or Fer-1 (10µM) co-treatment.

**Extended Data Figure 9.**
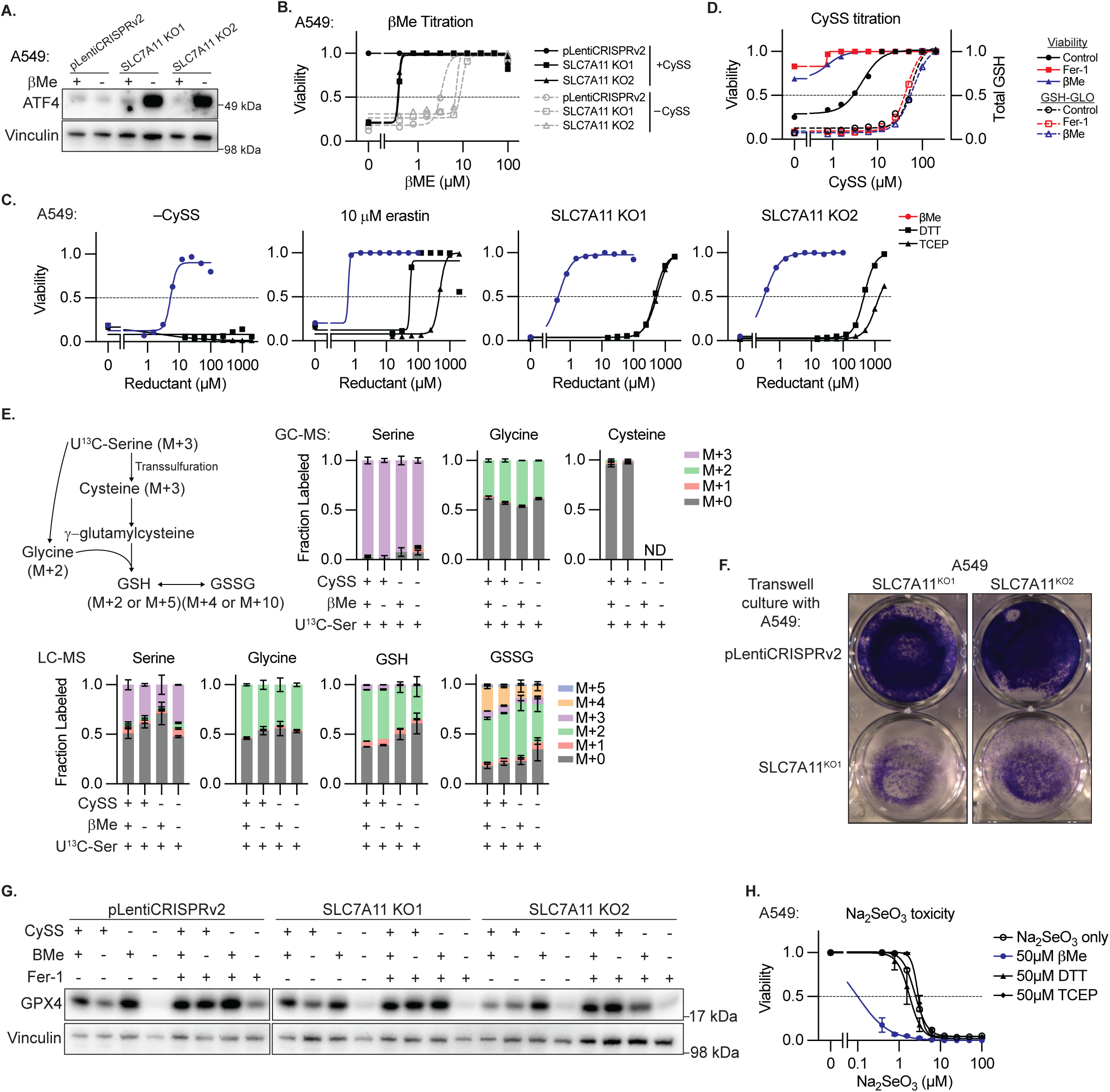
Data Supporting Figure 5. **A**, Immunoblot from lysates of A549 cells or SLC7A11 knockout clones for indicated proteins upon treatment with βMe (50μM). Inferred MW of band indicated. **B**, Viability (ATP) of cells from C cultured in the presence or absence of cystine (CySS) with indicated concentration of supplemented βMe for 3 days. Data report mean of technical triplicates. **C**, Viability (ATP) of A549 cells in cystine-free media or upon erastin treatment (10μM), or SLC7A11 knockout clones in standard conditions, treated with the indicated concentrations of reducing agents βMe, dithiothreitol (DTT), or Tris(2-carboxyethyl)phosphine (TCEP), 3 days. Data are the mean of two technical replicates. **D**, Viability (ATP, solid lines) and GSH content (dashed lines) of A549 cells cultured at indicated CySS concentration for 3 days. Data report mean of two technical replicates. **E**, Upper left, schematic delineating the contribution of serine-derived carbon to cysteine, glycine, GSH, or GSSG. Right, isotopic labeling of the indicated metabolites derived from A549 cells cultured in U-^13^C-serine (286 μM) as the only source of serine (top, GC-MS detection; bottom, LC-MS detection), in the presence or absence of CySS or βMe (50 μM) for 24 hr. ND = not detected. **F**, Crystal violet staining of A549 SLC7A11 knockout clones from Figure 5C upon transwell culture with A549 cells expressing pLentiCRISPRv2 or an SLC7A11 knockout clone, 3 days. Data are from one experiment. **G**, Immunoblot from lysates of A549 cells or SLC7A11 knockout clones for indicated proteins in the presence or absence of CySS, βMe (50 μM) of Fer-1 (10 μM) for 2 days. Inferred MW of band indicated. Data are from one experiment. **H**, Viability (ATP) of A549 cells upon treatment with Na2SeO3 and the indicated concentrations of reducing agents βMe, DTT, or TCEP, 3 days. Data are the mean ± SEM of n=3 biological replicates.

**Extended Data Figure 10.**
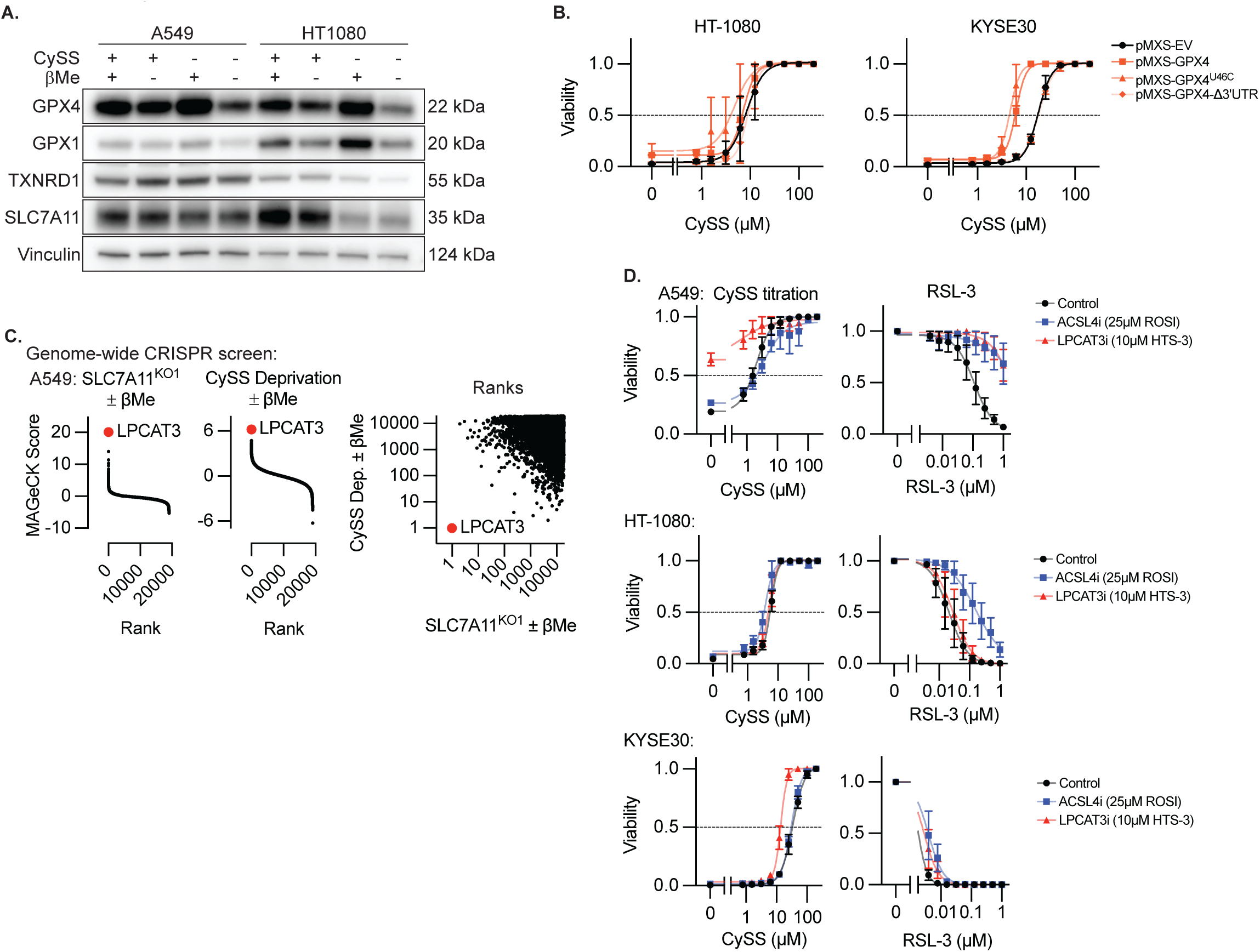
Cystine-deprivation sensitive cell lines that fail to be rescued by reducing agents undergo ferroptosis that is largely independent of ACSL4, and LPCAT3. **A**, Immunoblot from lysates of A549 cells or HT-1080 cells for indicated proteins in the presence or absence of CySS or βMe (50μM). Inferred MW of band indicated. **B**, Viability (ATP) of HT-1080 or KYSE30 cells expressing the indicated GPX4 cDNAs and selenocysteine (U46C) or SEICS (Δ3’UTR) mutants, cultured in indicated CySS conditions, 3 days. Data report mean ± SEM of n=3 biological replicates. **C**, Model-based Analysis of Genome-scale CRISPR-Cas9 Knockout (MAGeCK) score from genome-wide CRISPR screens of A549 SLC7A11 knockout cells or A549 cells cultured in the absence of cystine comparing culture in the absence or presence of βMe (50μM). LPCAT3 (red dot, indicated) is the top ranked gene in both conditions for promoting survival in the absence of βMe. Data are from a single screen. **D**, Viability (ATP) of indicated cell lines cultured in indicated CySS conditions or RSL-3 dose, in combination with inhibitors of ACSL4 (ROSI, 25μM) or LPCAT3 (HTS-3, 10μM), 3 days. Data report mean ± SEM of n=3 biological replicates.

## MATERIALS AND METHODS

### Reagents

Reagents were obtained from the following sources.

#### Antibodies

SLC7A11 (12691), Vinculin (4650), ATF4 (11815), RPS6 (2217) from Cell Signaling Technology. GPX4 (ab125066) and GPX1 (ab22604) from abcam. TXNRD1 (SC-28321), GCLC (SC-166345) from Santa Cruz Goat Anti-mouse IgG (H+L), HRP (Invitrogen; 62-6520) Goat Anti-rabbit IgG (H+L), HRP (Invitrogen; 31460).

#### Compounds and Chemical Reagents

RPMI (11875093, ThermoFisher); PrimeStar HS DNA Polymerase Premix (R040A, Takara); RNeasy Plus Mini Kit (74136), Qiaprep Spin Miniprep Kit (27106), and QIAquick Gel Extraction Kit (28706) from Qiagen. L-buthionine sulfoximine (B2515), L-buthionine sulfoxide (SC-207795, Santa Cruz), L-methionine sulfoximine (AC227202500, Fisher Scientific), L-methionine sulfoxide (36255-1, Cayman Chemical), Cystine (C7602, Sigma), Erastin (S7242, Selleckchem), RSL-3 (HY-100218A, MedChemExpress), beta-mercaptoethanol (NC9604560, Fisher Scientific), ferrostatin-1 (SML0583, Sigma), cycloheximide (14126, Cayman), eprenetapopt (gift from Aprea Therapeutics), DMSO (3176, Cell Signaling Technologies), sodium selenite (214485, Sigma), doxycycline (J60579.22, LifeSciences), ML-162 (HY-100002, MedChemExpress) ML-210 (HY-100003, MedChemExpress), DTT (D0632, Sigma), TCEP (C4706, Sigma), U-13C-serine (CLM-1574-H, Cambridge Isotopes), Rosiglitazone (71740, Cayman), (R)-HTS-3 (37563, Cayman), GSH-Glo (Promega V6911), Cell Titer Glo (Promega G7573), Cell Death Inducer Compound Library (35093, Cayman), Transwell culture apparatus (CLS3464, Sigma).

#### Cell Lines

5637, A-172, 786-O, A-253, A-375, A549, ACHN, AGS, AN3 CA, AsPC-1, BT-20, BT-549, BxPC-3, CAL 27, Calu-1, CAMA-1, DBTRG-05MG, Detroit 562, DoTc2 4510, DU 145, FaDu, HCC70, HCC827, HCT-15, HEC-1-A, HEC-1-B, Hep G2, Hs 578T, HT-1080, J82, LN-229, LNCaP, MDA-MB-231, MIA PaCa-2, NCI-H2052, NCI-H647, NCI-N87, PC-3, SH-SY5Y, SK-ES-1, SK-MEL-2, SK-OV-3, SNU-387, SNU-423, TT, U-2 OS, U-87 MG, UM-UC-3, DLD-1, HT-29, HCT-116, SW-620, MDA-MB-468, MDA-MB-453, T47D, ZR-75-1, MCF-7, HeLa, AGS, ACHN, A-498, HCC1438, NCI-H2030, SK-MES-1, PANC-1, PATU-8988T, RPE-1 from American Type Culture Collection; 647-V, BEN, CAL-33, EFM19, KYSE-30, OE-33, SK-MEL-30 from DSMZ, KP-2, OUMS-23, TIG-101, TIG-102, TIG-121, TIG-3S, TIG-120, TIG-107 from JCRB; SUDHL-4, SUDHL-1, SUDHL-5, DEL, OCILY-8, NALM-6, Jurkat, MOLT-4, SUDHL-7, NB-4, SKM-1, THP-1, KASUMI-1, MOLM-13, K-562, 293FT from the Broad Institute CCLE; HL-60, RAJI, and HMEC-E were provided by R. Weinberg (MIT). HFF isolation was previously described.^66^

#### Plasmids

sgRNAs/crRNAs were expressed in pLentiCRISPR-v2 (Addgene 52961) or pRDA-550 (Addgene 203398) and sequences are provided in Supplemental Table 7. cDNA vectors for exogenous expression of GPX4, SLC7A11, GCLC, or TXNRD1 and their mutants were synthesized as dsDNA (Twist Biosciences) and cloned into pMXS-IRES-BLAST (CellBioLabs), or pCW57.1 (Addgene 41393). Resulting plasmids are in the process of being deposited at Addgene.

### Cell culture

Cells were cultured in RPMI (11875093, ThermoFisher) supplemented with 10% fetal bovine serum (Peak Serum) and 1% penicillin and streptomycin (15140122, ThermoFisher), passaged using 0.05% Trypsin (15090046, ThermoFisher), diluted in PBS, and maintained at 5% CO2. Cell lines were tested for mycoplasma by polymerase chain reaction (PCR), and the authenticity of a subset of cell lines was verified by short tandem repeat (STR) profiling. Single-cell knockout clones are obtained by infecting parental cells with pLentiCRISPR-v2 into which specific sgRNAs were cloned. After selection, cells are plated in 96 wells in serial dilution and single clones are screened by immunoblotting. For experiments in which amino acids were removed from the cell culture media, base media lacking amino acids (R9010-01, US Biological) and dialyzed FBS was used. Dialyzed FBS was prepared by transferring FBS to dialysis tube (Fisher Scientific; molecular weight cut-off: 3,500 Dalton), and twice dialyzed against 4 L of sterile PBS with stirring for 6-12 hours. Dialyzed FBS was filtered through a 0.22 µm PES filter and stored at -20C.

### Phenotypic modulatory profiling screen

A panel of 100 human cancer cell lines was screened using a phenotypic modulatory profiling approach. Cell lines were thawed and maintained in RPMI-1640 supplemented with 10% fetal bovine serum and 1% penicillin–streptomycin for at least 7 days prior to screening. For adherent cell lines, 2,500 cells per well were plated in white-walled 96-well plates in 100 µL complete medium and allowed to adhere overnight. Twenty-four hours after plating, media were aspirated and replaced with 50 µL of medium containing 2× concentrations of rescue modulators (ferrostatin-1 (Fer-1), β-mercaptoethanol (βMe), or DMSO vehicle control). Subsequently, 50 µL of 2× compound solutions were added to achieve final 1× concentrations in a log₂-spaced 9-point dose–response series. Top concentrations were as follows: RSL3 (1 µM), erastin (10 µM), BSO (1 mM), and eprenetapopt (100 µM). For cystine titration experiments, cells were cultured in RPMI containing 10% dialyzed fetal bovine serum. Rescue modulators (2×) were added in cystine-free medium (50 µL), followed by addition of 50 µL of 2× cystine solutions to generate a titration series (maximum final concentration 200 µM). For suspension cell lines, 5,000 cells per well were plated in 50 µL medium in white-walled 96-well plates. Rescue modulators were added as 25 µL of 4× stock solutions, followed by 25 µL of 4× compound stocks to achieve final 1× concentrations in a 100 µL total volume. For cystine titration in suspension lines, cells were plated in 50 µL cystine-free medium, followed by addition of 25 µL 4× rescue modulators in cystine-free medium and 25 µL 4× cystine titration solutions. Cells were exposed to compounds or cystine titration conditions for 72 hours. Cell viability was assessed by addition of 40 µL CellTiter-Glo reagent directly to each well. Plates were shaken for 10 minutes at room temperature and luminescence was measured using a SpectraMax M3 plate reader (Molecular Devices). Viability was calculated by normalizing luminescence values to vehicle-treated controls for drug response experiments or to 200 µM cystine for cystine titration experiments within each rescue condition (DMSO, Fer-1, or βMe). Maximum viability was set to 1 for each condition. Dose–response data were summarized by calculating the area under the curve (AUC). For each condition, AUC was computed by summing normalized viability values across the 9 dose points, yielding a value ranging from 1 to 10, with higher values indicating greater overall viability across the dose range. To enable comparison across treatment groups, AUC values were standardized within each treatment group. Specifically, normalized AUCs were calculated by z-score transformation using the mean and standard deviation of the corresponding DMSO control AUCs for that treatment group. This normalization controlled for baseline viability differences and enabled cross-condition comparisons.

### Cloning of sgRNA and crRNA libraries

CRISPR libraries were designed, synthesized, and cloned as previously described with modifications. ^67^ Guide RNAs were designed using CRISPick to generate a Cas9 knockout (KO) library containing 10 sgRNAs per gene. In parallel, tiling libraries were generated targeting AIFM2, GCLC, GCLM, GPX4, GSR, GSS, and SLC7A11 using all possible SpCas9 sgRNAs (PAM: NGG) and enAsCas12a crRNAs across coding regions annotated from the human GRCh38 reference genome (Ensembl v1.115). Libraries included intragenic and non-targeting negative controls (50 each for ferroptosis Cas9 KO and enAsCas12a tiling libraries; 100 for ferroptosis Cas9 tiling libraries) and 10 sgRNAs per essential gene (DBR1, EEF2, GAPDH, KIF11, PCNA, PLK1, POLR2B, PSMB1, RPA3, and RPL3) for the Cas9 KO library. Final library sizes were 270 sgRNAs (Cas9 KO), 1,456 guides (enAsCas12a tiling), and 3,209 guides (SpCas9 tiling). BsmBI recognition sites, cloning overhangs, and subpool primer binding sites were appended to each 20-nt sgRNA or 23-nt crRNA sequence, resulting in 83-nt (SpCas9) and 85-nt (enAsCas12a) oligonucleotides. Oligonucleotide structures were as follows: SpCas9 libraries: 5′–[forward subpool primer]–CGTCTCACACCG–[20-nt sgRNA]–GTTTCGAGACG–[reverse complement of reverse subpool primer]–3′ and enAsCas12a libraries: 5′–[forward subpool primer]–CGTCTCAAGAT–[23-nt crRNA]–GAATCGAGACG–[reverse complement of reverse subpool primer]–3′. Oligonucleotide pools were synthesized by Twist Bioscience.

### Library amplification and cloning

Individual subpools were PCR-amplified in at least four parallel 50 µL reactions containing 25 µL NEBNext High-Fidelity 2× PCR Master Mix (M0541L, New England Biolabs), 1 µL oligonucleotide pool (5 ng), 2.5 µL each of 10 µM forward and reverse primers, and nuclease-free water to volume. PCR cycling conditions were: 98°C for 30 s; 24 cycles of 98°C for 10 s, 53°C for 30 s, 72°C for 30 s; followed by a final extension at 72°C for 2 min. Amplicons were pooled and purified using the Monarch PCR & DNA Cleanup Kit (T1130L, New England Biolabs). Purified products were cloned by Golden Gate assembly into BsmBI-v2–digested pLentiCRISPRv2 (Cas9 libraries) or pRDA-550 (enAsCas12a libraries) using the NEBridge Golden Gate Assembly Kit (E1602L, New England Biolabs). Assembly reactions were cycled 65 times according to the manufacturer’s protocol. Ligation products were isopropanol-precipitated and electroporated into Stbl4 electrocompetent cells (Invitrogen) using a Gene Pulser Xcell Electroporation System (Bio-Rad). Cells were recovered for 90 min at 30°C in SOC medium and cultured for 16 h at 30°C in LB broth supplemented with 100 µg/mL carbenicillin. Library representation was estimated by serial dilution and plating on LB agar plates in parallel. Plasmid DNA was isolated using the HiSpeed Plasmid Maxi Kit (Qiagen). Library composition and guide representation were confirmed by next-generation sequencing.

### Target identification screens

HT-1080 cells were transduced with pooled lentiviral libraries at a multiplicity of infection (MOI) of 0.3–0.5 to favor single-guide integration. A total of 12 × 10^6^ cells were plated in 12-well plates (3 × 10^6^ cells per well) and transduced in the presence of 8 µg/mL polybrene. Plates were centrifuged at 1,200 × g for 1 hour at 37°C to enhance transduction efficiency. Twenty-four hours after transduction, cells were trypsinized, pooled, and replated in 15-cm dishes for recovery. Puromycin selection (2 µg/mL) was applied for 72 hours to enrich for transduced cells. Following selection, cells were expanded for an additional 3 days prior to treatment initiation. For screening, cells were split into drug treatment conditions in technical duplicate while maintaining >1,000× library representation at all times. For each condition, 2.3 × 10^6^ cells were plated per 15-cm dish. A baseline sample was collected at the time of treatment initiation. Drug treatments were as follows: RSL3 (1 µM), erastin (5 µM), BSO (100 µM), ML-162 (1 µM), ML-210 (5 µM) and eprenetapopt (50 µM). For the SpCas9 tiling screen, one 15-cm plate per replicate was used per condition; for the enAsCas12a tiling screen, two 15-cm plates per replicate were used per condition to maintain representation. Cells were passaged, counted, and re-dosed every 3 days for a total of 15 days. At the endpoint, cells were harvested and genomic DNA was extracted for crRNA/sgRNA quantification by next-generation sequencing.

### Targeted amplicon sequencing

To characterize mutations generated by validation crRNAs using enAsCas12a, HT-1080 cells were transduced with individual crRNAs cloned into the pRDA-550 vector. Briefly, 0.5 × 10^6 cells were transduced and selected with puromycin for 3 days to enrich for transduced cells. Following selection, cells were expanded for an additional 6 days prior to drug treatment. Cells were then exposed to ferroptosis-inducing compounds for 15 days, including RSL3 (1 µM), erastin (5 µM), or BSO (100 µM). Following treatment, genomic DNA was isolated from surviving cells as well as from untreated HT-1080 parental cells. Regions flanking the crRNA target sites were PCR-amplified using locus-specific primers and PrimeSTAR HS DNA Polymerase (Takara, R010A). PCR amplicons were submitted for next-generation sequencing (Amplicon-EZ; 150–500 bp, Genewiz). Sequencing reads were analyzed using CRISPResso2 to quantify insertion–deletion (indel) frequencies and mutation profiles at target loci.

### G enome-wide CRISPR screens

Genome-wide resistance screens were performed using the Human Activity-Optimized CRISPR Knockout Library (Addgene pooled library #1000000100). A549 parental cells or an isogenic SLC7A11 knockout clone were transduced with pooled lentivirus at a multiplicity of infection (MOI) of approximately 0.3–0.5 to favor single-guide integration. 72 × 10^6^ cells were plated in 12-well plates (3 × 10^6^ cells per well) to maintain ∼100× library representation at the time of transduction. Following puromycin selection (Parental: 2 µg/mL, SLC7A11 KO: 4 µg/mL), cells were expanded in complete medium supplemented with 50 µM β-Meto maintain viability. To identify genes associated with resistance to βMe withdrawal, ∼18 × 10^6^ A549 parental cells per condition (2.3 × 10^6^ cells per 15-cm plate; ∼100× representation) were plated in medium containing dialyzed fetal bovine serum under cystine-deprived conditions in the presence or absence of 50 µM βMe. In parallel, A549 SLC7A11 knockout cells were plated in standard serum-containing medium with or without 50 µM βMe. Cells were passaged and re-plated every 3 days while maintaining library representation for a total of 15 days. At the experimental endpoint, cells were harvested and genomic DNA was extracted for sgRNA amplification and quantification by next-generation sequencing.

### Immunobloting

Cells are washed with cold phosphate-buffered saline and lysed in RIPA lysis buffer [50 mM tris (pH 7.4), 150 mM NaCl, 1% NP-40, 0.1% sodium deoxycholate, 0.1% SDS, and 2 mM EDTA] with a protease inhibitor cocktail (Sigma-Aldrich, 5892791001). Cell lysates are collected after 10-min incubation on ice and centrifuged at 4°C at 21,000g for 10 min. Protein concentration is quantified using a bicinchoninic acid (BCA) protein assay kit (Pierce 23225). Eight micrograms protein is loaded with loading buffer into Bolt 4 to 12% bis-tris polyacrylamide gels (Thermo Fisher Scientific, NW04125BOX) with protein ladder (Thermo Fisher Scientific, LC5925), electrophoresed in running buffer (Thermo Fisher Scientific, B0002) at 110 V for 1 hours, and transferred in transfer buffer [N-cyclohexyl-3-aminopropanesulfonic acid (2.2 g liter−1), NaOH (0.45 g liter−1), and 10% ethanol] to a polyvinylidene difluoride membrane (Millipore, IPVH00010) at 75 V for 3.5 hours. Membranes are blocked with 5% bovine serum albumin (BSA), incubated with indicated antibody for 1-2hrs at room temperature, washed three times with TBS-T (tris buffered saline, 0.1% Tween-20), incubated with secondary antibody (1:10,000) for 1 hour at room temperature, washed three times with TBS-T, and developed using ECL substrate (Thermo Fisher Scientific, PI-32106) on a CCD-camera-based detector. Each experiment is replicated three times, representing figures are shown in the results.

### Metabolomics

#### GC/MS

1 x 106 A549 cells in a 6-well plate were placed in the indicated conditions and treated with 100 μM U-13C-serine tracer in media lacking serine for 24 hours. Labeled cells were washed with cold 0.9% saline, and 500 µL of cold 80% methanol containing 1 µg/mL norvaline (Sigma) added. Cell extracts were scraped into 1.5 mL pre-chilled Eppendorf tubes, vigorously vortexed at 4C for 10 minutes, and centrifuged at 14.8 x 103 RPM (21.1 x 103 g), 4C for 10 minutes. The supernatant was dried using a Speedvac. Dried metabolites were treated with 20 µL of 20 mg/mL of methoxyamine hydrochloride (MOX; in pyridine; Supelco) 1 hour at 37C (1st derivatization), and silylated either by 20µL TMS (N-methyl-N-(trimethylsilyl)trifluoroacetamide; Supelco) 1 hour at 37C (2nd derivatization). Resulting derivatized samples were transferred to a Target DP vial (Thermo Scientific; TFC4000-11) and sealed with a screw cap (Agilent; 5185-5820). Samples were run on an Agilent Technologies 5977B mass spectrometer (MS) coupled with Agilent Technologies Intuvo 8000 gas chromatographic system (GC). Software: 2. Intuvo 9000 GC Firmware; version 2.4.1.10; for GC-MS (Agilent), as described.^68^

#### LC/MS

1 x 106 A549 cells in a 6-well plate were placed in the indicated conditions and treated with 100 μM U-13C-serine tracer in media lacking serine for 24 hours. Labeled cells were washed with cold 0.9% saline, scraped, transferred into microcentrifuge tubes, and pelleted. Cell pellets were submitted to the NYU Metabolomics Laboratory, which extracted metabolites in 80% methanol, and ran reconstituted samples on a Thermo Scientific Q Exactive Hybrid Quadrupole-Orbitrap Mass Spectrometer.

### Animal Use

Tumors were initiated into 4-8 week old female NCr athymic nude (Taconic), NSG mice or Balb/c mice (Jackson Labs) in the right flank by implanting 5 × 10^6^ cells in 50% matrigel in a total volume of 100 μL, except for genetic screens in which case 5 × 10^6^ cells were implanted in each flank. Expression of DOX-controlled genes was induced by switching mice onto doxycycline chow (1000 mg/kg) upon formation of 100 mm^3^ tumors. Tumor volume was assessed using caliper measurements and volume was calculated using (L ×W^2^)/2. For genetic screens, tumors were harvested after 14 days (HT-1080) or 21 days (A549), and DNA was isolated from homogenized tumors for downstream processing. Where indicated, BSO was provided in the drinking water for the indicated durations. All experiments involving mice were carried out with approval from the Committee for Animal Care and under supervision of the Department of Comparative at NYU Langone Medical Center.

### Data Analysis

Measurements were taken from distinct samples. Statistical tests comparing two groups were performed using a two-sided heteroscedastic student’s t test. Statistical comparisons between experimental groups in animal studies were performed using a two-tailed Mann–Whitney U test (nonparametric rank-based test).

## ACKNOWLEDGEMENTS

We thank Possemato and Pacold lab members for helpful comments. pLentiCRISPR v2 was a gift from Feng Zhang. **Funding:** National Institutes of Health R01CA286141 (RP), R01GM132491 (RP), Cancer Center Support Grant P30CA016087 (Kimmelman).

## Author contributions

Conceptualization and Methodology (Figure 1): KMF, RP, MEP; Conceptualization and Methodology (Figures 2-5): KMF, RP; Investigation: KMF, AA, BA, MGP, GF, KW, NJC, EMT; Funding acquisition, resources, and supervision: RP, MEP; Manuscript preparation – initial draft: RP, KMF. All authors reviewed and approved the manuscript.

## Data and Resource Availability

All data supporting the findings of this study are available within the paper and its Supplementary Information. All materials used in this study are readily available. Modified cell lines can be obtained from the corresponding author upon request.

